# Cataract-prone variants of γD-crystallin populate a conformation with a partially unfolded N-terminal domain under native conditions

**DOI:** 10.1101/2024.06.01.596973

**Authors:** Sara Volz, Jadyn R. Malone, Alex J. Guseman, Angela M. Gronenborn, Susan Marqusee

## Abstract

Human γD-crystallin, a monomeric protein abundant in the eye lens nucleus, must remain stably folded for an individual’s entire lifetime to avoid aggregation and protein deposition-associated cataract formation. γD-crystallin contains two homologous domains, an N-terminal domain (NTD) and a C-terminal domain (CTD), which interact via a hydrophobic interface. A number of familial mutations in the gamma crystallin gene are linked to congenital early-onset cataract, most of which result in amino acid changes in the NTD. Several of these, such as V75D and W42R, are known to populate intermediates that, under partially denaturing conditions, possess a natively folded CTD and a completely unfolded NTD, with studies on W42R showing further evidence for a minor population of an intermediate under native conditions. We employed hydrogen-deuterium exchange mass spectrometry (HDX-MS) to probe the structural and energetic features of variants of γD-crystallin under both native and partially denaturing conditions. For V75D and W42R, we identify a species under native conditions that retains partial structure in the NTD and is structurally and energetically distinct from the intermediate populated under partially denaturing conditions. Residues at the NTD-CTD interface play crucial roles in stabilizing this intermediate, and disruption of interface contacts either by amino acid substitution or partial denaturation permits direct observation of two intermediates at the same time. The newly identified intermediate exposes hydrophobic amino acids that are buried in both the folded full-length protein and in the protein’s stable isolated domains. Such non-native exposure of a hydrophobic patch may play an important role in cataract formation.

**Significance Statement:** Human γD-crystallin, which plays a structural role in the eye lens, is a long-lived protein that must remain folded for an individual’s entire lifetime to avoid aggregation and protein deposition - associated cataract formation. By using hydrogen-deuterium exchange mass spectrometry, we demonstrate that two cataract-associated variants of γD-crystallin populate an intermediate with partial structure along the interface between its two domains under native conditions. In these intermediates, hydrophobic amino acids that are normally buried in the N-terminal domain’s native folded structure become exposed, possibly leading to aggregation and cataract formation. Our findings illustrate the importance of studying a protein’s energy landscapes under conditions that are close to physiological.

## Introduction

The lens of the eye is a unique and highly specialized organ subject to specific biophysical constraints. It must be transparent to prevent light scattering or absorption, it must possess a high refractive index to tightly focus light onto the retina, and its focal length must be adjustable to permit focusing over a range of distances (1, 2). To achieve these optical properties, 90% of the protein mass in fully differentiated lens fiber cells consists of soluble, densely packed crystallin proteins at concentrations exceeding 400 mg/ml (3). During lens maturation, all cellular structures that contribute to light scattering are eliminated, including nuclei and organelles, effectively stopping protein turnover in the mature lens (4, 5). Therefore, to maintain faithful functioning of the eye lens, crystallins must remain soluble and stably folded for an individual’s entire lifetime. Alas, over years, crystallins are exposed to environmental and chemical assaults, resulting in damage that can initiate aggregation and precipitation. Subsequent opacification of the lens, as seen in age-related cataract, is a condition that affects more than 20 million people and is the leading cause of blindness worldwide (6).

Crystallins are subdivided into three families: α-, β-, and γ-crystallins. α-crystallins are small heat shock proteins that assemble into oligomeric structures, perform chaperone functions, and assist in homeostasis of β- and γ-crystallins (7). β- and γ-crystallins, which exist as dimers/oligomers and monomers, respectively, are small, stable proteins with high solubility that play structural roles in conferring the optical properties of the eye lens (8). Human γD-crystallin (γDC) is the second-most abundant γ-crystallin found in the lens nucleus, comprising ∼11% by mass of the total protein in young human lenses (9). γDC is relatively small (∼21 kDa) with two homologous β-sheet domains, an N-terminal domain (NTD) and a C-terminal domain (CTD), each comprised of two Greek key motifs (10). The CTD and NTD are connected by a short linker and interact via a nonpolar interface (Figure 1A) (11).

**Figure 1.**
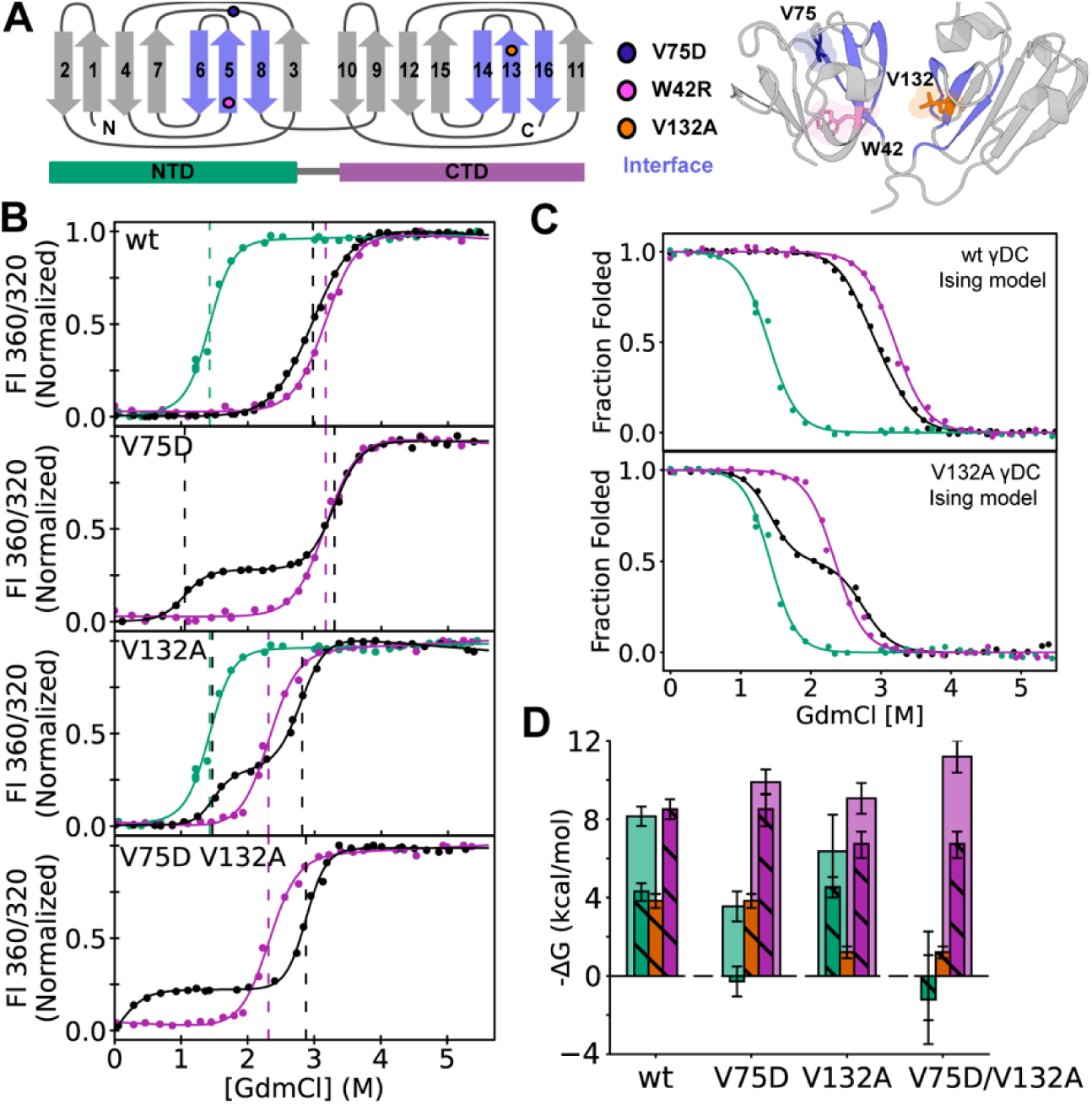
Equilibrium denaturation experiments of humanγD-crystallin variants. A) Topological map and crystal structure (1HK0 (10)) of γDC, with Greek key beta strand architecture, NTD and CTD, and three residues of interest indicated. β-strands containing interface interactions between the two domains are highlighted in blue. B) Two- and three-state fits to tryptophan fluorescence data of full-length constructs and their corresponding isolated individual domains (γDC_NTD_, green; γDC_CTD_; purple; full-length γDC, black). C_M_ values are indicated with dotted lines to draw the eye. C) Global Ising fits from analysis of the same data for V132A and wild-type (γDC_NTD_, green; γDC_CTD_, purple; full-length γDC, black). D) Graphical estimation of the relative energetic contributions of each domain’s intrinsic stability and interfacial interaction energy to each transition (NTD, green; CTD, purple; interface, orange). Total transitions (solid bars)—that is, transitions calculated irrespective of dissection into component intrinsic and interaction energies—are taken from three-state fits, except in wild-type, where a three-state fit could not be obtained and the Ising interface and intrinsic γDC_NTD_ energies were summed to estimate the total NTD transition. Intrinsic and interfacial energies (striped bars) are based on values from Ising fits. In the case of constructs containing V75D where Ising fits could not be obtained, the corresponding interaction energy in wild-type or V132A was subtracted from the construct’s three-state NTD transition to estimate the intrinsic energy of V75D γDC_NTD_ and V75D/V132A γDC_NTD_, respectively, under the assumption that the V75D mutation does not impact the interaction energy. Error bars represent standard error of the fit. Parameters for all fits may be found in Table 2. All data were collected at 25°C in PBS pH 7.0, 5 mM DTT.

Despite the high thermodynamic and kinetic stability of γDC, protein damage, by UV (12), deamidation (13), oxidation (14), or other modifications, has been proposed to result in aggregation and cataract (15). Furthermore, mutations in the γD-crystallin gene have been linked to early-onset or congenital cataract, including R14C (16), P23T (11, 17), W42R (18), R58H (19), G60C (20), V75D (21), R76S (22), and I81M (23), among others. Interestingly, for most amino acid changes, including V75D (24) and W42R (25), no significant structural changes are observed, suggesting that differences in folding/unfolding behavior (that is, in the energy landscape) are likely the source of the aggregation-prone behavior.

Upon chemical denaturation, wild-type γDC populates a stable intermediate best characterized as a fully unfolded NTD and a fully folded CTD (26–30). Isolated N-terminal and C-terminal domain fragments of γDC (hereafter termed γDC_NTD_ and γDC_CTD_) are stable and well-folded in isolation. γDC_NTD_ is less stable than γDC_CTD_ despite the near-equivalence of free energies for the two unfolding transitions in the full-length protein (27, 31). Therefore, the presence of the CTD stabilizes the NTD in the full-length protein, most likely due to amino acids in the NTD (C41, M43, F56, R79, I81, and P82) that face the opposing CTD surface (comprising residues V132, G141, Q143, L145, R167, and V169) (11). Past work has estimated this contributes 4 kcal/mol (31), and destabilization of these predominantly hydrophobic interactions, as in the CTD variant V132A, results in destabilization of the NTD unfolding transition (30).

Destabilization of the NTD directly, as in V75D, results in population of an intermediate across a broad range of denaturant concentrations (24). This intermediate has been structurally characterized by both NMR and SAXS and is comprised of a fully unfolded NTD and a fully folded CTD (32). These studies raise the tantalizing possibility that this partially unfolded species might serve as a specific nucleus for aggregation and/or cataract formation. Furthermore, destabilization of the NTD in the variant W42R permits observation of a minor species with unfolding in the NTD under native conditions (25).

These partially structured states accessible under non-native conditions may not be identical to the conformations accessed under physiological or native conditions. We therefore used hydrogen–deuterium exchange coupled with mass spectrometry (HDX-MS) to investigate the structural and energetic characteristics of γDC’s equilibrium intermediate(s) under both native and denaturing conditions. While most standard techniques for probing protein stability, such as circular dichroism or fluorescence, are unable to detect minor conformations at low population, HDX-MS has the unique capability to both identify and structurally characterize such rare states.

HDX-MS monitors the exchange of labile hydrogens with solvent deuterons (33). For backbone amides, this exchange requires accessibility to the deuterated solvent and/or transient hydrogen-bond opening. Therefore, fast HDX rates correlate to exposed and flexible regions in the protein, while slow rates are associated with amide hydrogens that are buried within the protein and/or involved in stable hydrogen bonds. The relationship between the observed rate of HDX can be analyzed using the Linderstrøm-Lang model (34) to determine either the energetics or kinetics for the opening event that exposes the amide. (35). HDX-MS is therefore an ideal tool to characterize systems like γDC, a protein for which potential sub-global unfolding events are particularly germane to function and misfunction.

We found that, for two cataract-associated variants of γDC with destabilizing amino acid changes in the NTD, the partially unfolded state detected under native conditions is not the intermediate populated under denaturing conditions, but rather a distinct species with a folded CTD and residual structure at the NTD/CTD interface. Moreover, disruption of hydrophobic interactions at the interface, either by introducing amino acid changes or by adding small amounts of denaturant, destabilizes this partially folded region, permitting direct observation of the two intermediates at once. Our data suggest that the newly identified intermediate is uniquely accessed from the native state and exposes a surface that normally is buried, both in the full-length structure and in the isolated folded NTD. It has been proposed that small populations of conformers that exhibit partial unfolding and are difficult to detect in solution may serve as the nucleus for aggregation, potentially resulting in cataract (25, 32, 36). Therefore, the newly discovered intermediate, which does not reside along the folding pathway, may represent the pivotal conformer that initiates the aggregation pathway ultimately resulting in cataract formation.

## Results

### Equilibrium unfolding/refolding of wild-type and variant proteins

HDX-MS permits detection of high-energy states. To estimate the expected stabilities of any such states identified by HDX-MS and compare them to known conformations, the relative contributions of the isolated NTD, the isolated CTD, and the NTD-CTD interface to the energetics of different crystallin variants need to be evaluated. Although many variants of γDC have already been characterized by chemically induced denaturation, so far all stability data were analyzed by standard two- or three-state equilibrium unfolding models, using linear extrapolations to zero denaturant (24–27, 30, 37). In order to calculate interaction energies precisely, and, more importantly, in a consistent manner, we repeated several γDC unfolding studies. We collected equilibrium unfolding curves for the full-length protein and isolated single-domains for several variants and analyzed the data using an Ising model (38, 39).

To measure global equilibrium unfolding, we used GdmCl-induced denaturation monitored by tryptophan fluorescence (Figure 1). γDC contains four tryptophan residues, two in the N-terminal domain (residues 42 and 68) and two in the C-terminal domain (residues 130 and 165). Since all four tryptophans are buried in the native structure and their fluorescence is quenched when γDC occupies the native state, intrinsic tryptophan fluorescence serves as an excellent reporter of unfolding in γDC (40, 41).

The variants we examined were wild-type γDC, V132A γDC, V75D γDC, V75D/V132A γDC, γDC with five amino acid changes in the CTD that reside in the NTD-facing side of the CTD (V132A/Q143A/L145A/M147A/V169A) (dInt-γDC), the isolated NTD (γDC_NTD_, residues 1-81), the isolated CTD (γDC_CTD_, residues 84-174), and the isolated CTD with V132A (V132A-γDC_CTD_) (Table 1). Data were first analyzed via either a two state (N ⇌ U) or three-state (N ⇌ I ⇌ U) linear-extrapolation model, resulting in parameters describing ΔG in the absence of denaturant, the dependence of ΔG upon denaturant (*m*-value) (42, 43). For V132A and wild type, data were subsequently analyzed using a 1D-Ising model (see below).

**Table 1.**
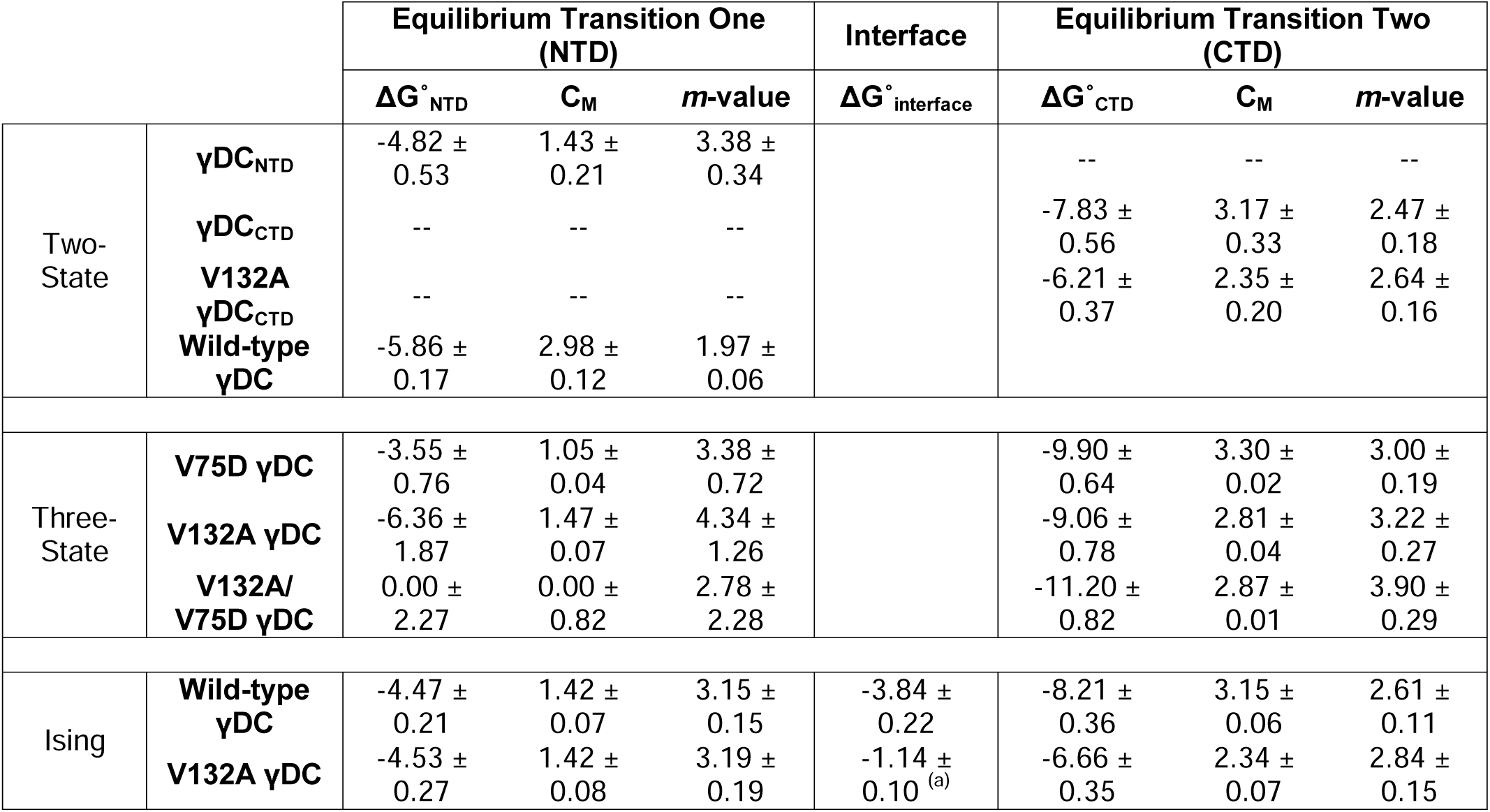
Equilibrium two-state, three-state, and global Ising fit parameters for allγDC variants. Error reported is standard error of the fit. ΔG° (folding) values are in kcal/mol; C_M_ values are in M GdmCl; *m*-values are in kcal/(mol* M GdmCl). Confidence intervals calculated from bootstrapped parameters for Ising fits can be found in Table S1. ^(a)^ The model for V132A includes a stabilizing interaction between the folded CTD and the unfolded NTD equal to that of the stabilizing interaction between the folded CTD and the folded NTD. A more complicated model permitting these two stabilizing interactions to vary independently of one another did not pass an F-test (p = 0.49) and so was not used.

All isolated domains studied (γDC_NTD_, γDC_CTD_, and V132A-γDC_CTD_) exhibit apparent cooperative two-state behavior, consistent with previously reported equilibrium data (27, 31) and were analyzed using the two state (N ⇌ U) model (Figure 1B) (Table 1). All full-length crystallin constructs studied, except for wild-type γDC, fit well to a three-state model (N ⇌ I ⇌ U). The un/folding isotherms for V75D, V132A, V75D/V132A, and dInt-γDC all show two transitions with a plateau, indicating the presence of an intermediate. For V132A, the transition to this intermediate begins at ∼1.5 M GdmCl, while in V75D the transition begins earlier, at around 1.0 M GdmCl (Figure 1B). Wild-type γDC has been previously observed to fit well to a three-state model (27, 31); however, under the conditions used here, the two transitions cannot be distinguished from one another, precluding a three-state fit. The resulting two-state fit, however, yields an unusually broad transition when compared to the isolated domains and is therefore considered to be unreliable, emphasizing the usefulness of an Ising model in dissecting the energetics of each transition (see below).

All full-length variants studied, whether containing amino acid changes in the NTD or the CTD, result in destabilization of the first unfolding transition relative to wild type, with V75D/V132A the most and V132A the least disruptive. Although V132A changes a residue in the CTD, it mostly impacts the first (NTD-unfolding) transition rather than the second, suggesting that removing a hydrophobic contact in the CTD region that faces the NTD negatively impacts the stability of the NTD in the context of the full-length γDC. In fact, the V75D/V132A variant is already in the unfolding transition at 0 M denaturant, indicative of the intrinsic instability of the NTD in V75D. These data are in agreement with similar studies, albeit collected under different conditions (e.g., temperature) (30).

The Ising model permits quantification of cooperativity between folding modules in a protein, and it has been successfully applied to repeat proteins wherein subsequent repeats interact via a defined interface (39, 44). Given measurements for the equilibrium unfolding transitions of both a full-length protein and its individual domains, this model can relate the extent of unfolding to the intrinsic ΔG of each domain and the interfacial ΔG upon association of the two domains. Therefore, we used a global fitting approach for a one-dimensional Ising model comprising a two-repeat heteropolymer as outlined previously (45) to analyze the folding transitions for the full-length γDC variants for which both isolated domain transitions were measured (wild type and V132A) (Figure 1C). Our results are summarized in Figure 1D.

The wild-type γDC data fits well to this 1D-Ising model (reduced sum of squared residuals (RSSR) = 4.11×10^-4^), with intrinsic folding free energies of -4.5 and -8.2 kcal/mol for the NTD and CTD domains, respectively. These ΔG values are in good agreement with those extracted from the equilibrium unfolding curves of isolated domains using two-state models. The interface contributes -3.8 kcal/mol in wild-type γDC, consistent with a previous estimate (31) (Table 1). We therefore conclude that the two domains are strongly energetically coupled, which has previously been suggested to explain the large kinetic barrier to unfolding of the NTD in the full-length protein (31).

For V132A, however, the equilibrium unfolding transition is not fit well by the standard 1D-Ising model, as evidenced by a non-random distribution of fit residuals (RSSR = 1.1×10^-3^, Figure S1). We note that the second unfolding transition (corresponding to unfolding of the CTD) is stabilized in the full-length V132A protein compared to the isolated domain (V132A-γDC_CTD_), even at concentrations of denaturant where the NTD is fully unfolded. Hence, we hypothesize that the presence of the unfolded NTD stabilizes the folded CTD in V132A γDC via an interaction that is not captured by the simplest version of the 1D-Ising model, possibly by providing hydrophobic interactions near position 132. Use of an interaction term for describing the stabilization of the folded V132A CTD by the NTD whether it is folded or unfolded (39) yields a significantly improved fit (RSSR = 5.12×10^-4^, Figure 1C). This interfacial energy was calculated to be -1.1 kcal/mol, representing a reduction in interfacial stability of 2.6 kcal/mol compared to wild type. An alternative, more complicated version of this model in which the stabilization of the CTD upon interaction with the unfolded NTD was used as a parameter distinct from the stabilization by the folded NTD, i.e. a model that uses two separate interaction terms, did not pass an F-test and was deemed not suitable (p = 0.49). In essence, our modified 1D-Ising model analysis of V132A γDC, shows, surprisingly, that interfacial stabilization can be conferred by the unfolded NTD, albeit to a lesser degree than by the folded domain in wild type. The strength of this interaction is likely similar in the V75D/V132A variant, as V75D does not impact the CTD transition. This implies that specific contacts between the two folded domains are lost upon removal of the hydrophobic sidechain of V132, but that stabilizing interfacial interactions in V132A are also mediated via the unfolded NTD. This is not the case for wild type: applying a folded-unfolded interaction term to wild type yields an interaction energy of ∼0 kcal/mol, suggesting that loss of specific contacts made by the valine side chain disrupts the interface (Table S1).

Additional changes to the CTD interface in dInt-γDC destabilize the second unfolding transition by 2.5 kcal/mol, but do not further disrupt the first unfolding transition compared to the single amino acid change in V132A (Figure S2A). Thus, these further changes to the CTD surface do not destabilize the NTD transition and instead only reduce the intrinsic stability of the CTD.

Finally, for technical reasons, we were unable to carry out a full Ising analysis for the V75D variant. The isolated V75D NTD (V75D-γDC_NTD_) proved refractory to purification and analysis, despite multiple attempts. Based on the above data, we predict that V75D-γDC_NTD_ is intrinsically unstable and that most of its stability in the full-length protein is provided by interfacial interactions. The low stability of the first transition of the V75D/V132A variant (in which stabilizing interactions with the CTD are reduced) also points to the intrinsic instability of V75D-γDC_NTD_.

### Continuous-labeling HDX-MS on variants of γD-crystallin

While the above equilibrium experiments and Ising analyses allow a quantitative determination of the stability of each domain and the interfacial coupling between them, they are unable to inform about the conformations associated with any partially unfolded states or to detect the presence of any low-occupancy (high-energy) states under native conditions. Therefore, we assessed the conformational ensembles of the γDC variants on a sub-global level using HDX-MS. We followed continuous hydrogen-deuterium exchange of wild-type and several variants of γDC over a time course spanning from 15 seconds to 72 hours, both in the presence and absence of denaturant (PBS pH 7.0, 5 mM DTT, 25 °C).

The kinetics of HDX are commonly analyzed using two kinetic regimes, EX1 and EX2. When monitored by mass spectrometry, these two kinetic limits can be distinguished by their mass spectra signatures: EX1 kinetics produce two distinct mass envelopes whose relative intensities change over the time of exchange without a shift in m/z, while EX2 kinetics result in the gradual increase in m/z over time of a unimodal mass envelope (46). For both regimes, the exchange process can be analyzed via the Linderstrøm-Lang model of HDX (see below, 36), where an amide hydrogen interconverts with rates *k_op_* and *k_cl_* between a closed, exchange-resistant state and an open, exchange-competent state.

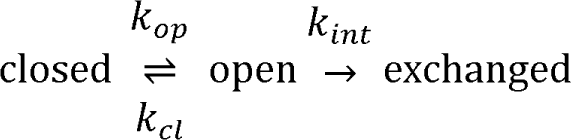

In this model, exchange in the open state occurs at a rate *k_int_*, the known chemical or intrinsic rate of exchange for a given amide hydrogen. When the rate of closing *k_cl_*is slow relative to intrinsic rate of exchange *k_int_* (*k_cl_* << *k_int_*), EX1 applies, and the observed exchange rate is equal to the rate of structural opening *k_op_*. Conversely, in the EX2 kinetic exchange limit, the rate of closing *k_cl_* is fast relative to the intrinsic rate of exchange *k_in_*_t_ (*k_cl_* >> *k_int_*). In this regime, the observed rate of hydrogen exchange is related to the free energy of the transition between the high-energy (open) and low-energy (closed) state; specifically, the observed rate constant of exchange can be expressed as *k_obs_* = (*k_op_* /*k_cl_)* \**k_int_* = K_op_\**k_int_*, with K_op_ the equilibrium constant for opening to the exchange-competent state. Thus, the free energy for opening is ΔG_op_ = -RT ln(*k_obs_/k_int_*). When exchange is monitored at the peptide level by mass spectrometry, the average intrinsic rate constant *k_int_* per peptide can be estimated based on the sequence of the individual peptide (47). Thus, if ΔG_unfolding_ of a structural transition is known, the expected HDX rates per peptide for opening associated with that transition can be estimated. Amide hydrogens in dynamic or flexible peptides can exchange more quickly than seen from such estimates. However, if EX2 exchange is significantly slower than estimated, a transition with a greater ΔG must be present.

Using aspergillopepsin and pepsin digestion of the proteins and a two-step quenching method, we obtained between 95-100% peptide coverage for all variants of γDC (see methods). This permitted interrogation of dynamics across all of γDC (average redundancy, coverage, and back exchange as well as other experimental parameters for all variants may be found in Table S2). In most variants, nearly all peptides exhibit a single isotopic envelope distribution whose centroid mass increases for increasing exchange times, demonstrating EX2 kinetic behavior, with a few notable exceptions (such as in the β1 strand, see below).

### γDC hydrogen-deuterium exchange is slow at the interface

For the V75D γDC variant, which populates an equilibrium intermediate in denaturant (Figure 1B), previous NMR studies in 4.2 M urea revealed an unfolded NTD and a native, folded CTD (32). HDX-MS data for the native state protein (0 M denaturant) or this intermediate (4.2 M urea) are summarized in Figure 2A-C. In 4.2 M urea, amides in NTD peptides exchange quickly, within 15 seconds, while amides in CTD peptides are slow to exchange, with exchange rates similar to those observed under native conditions (Figure S3). This is consistent with the structural results obtained by NMR. We term this intermediate observed under denaturing conditions the “exposed-interface intermediate.”

**Figure 2.**
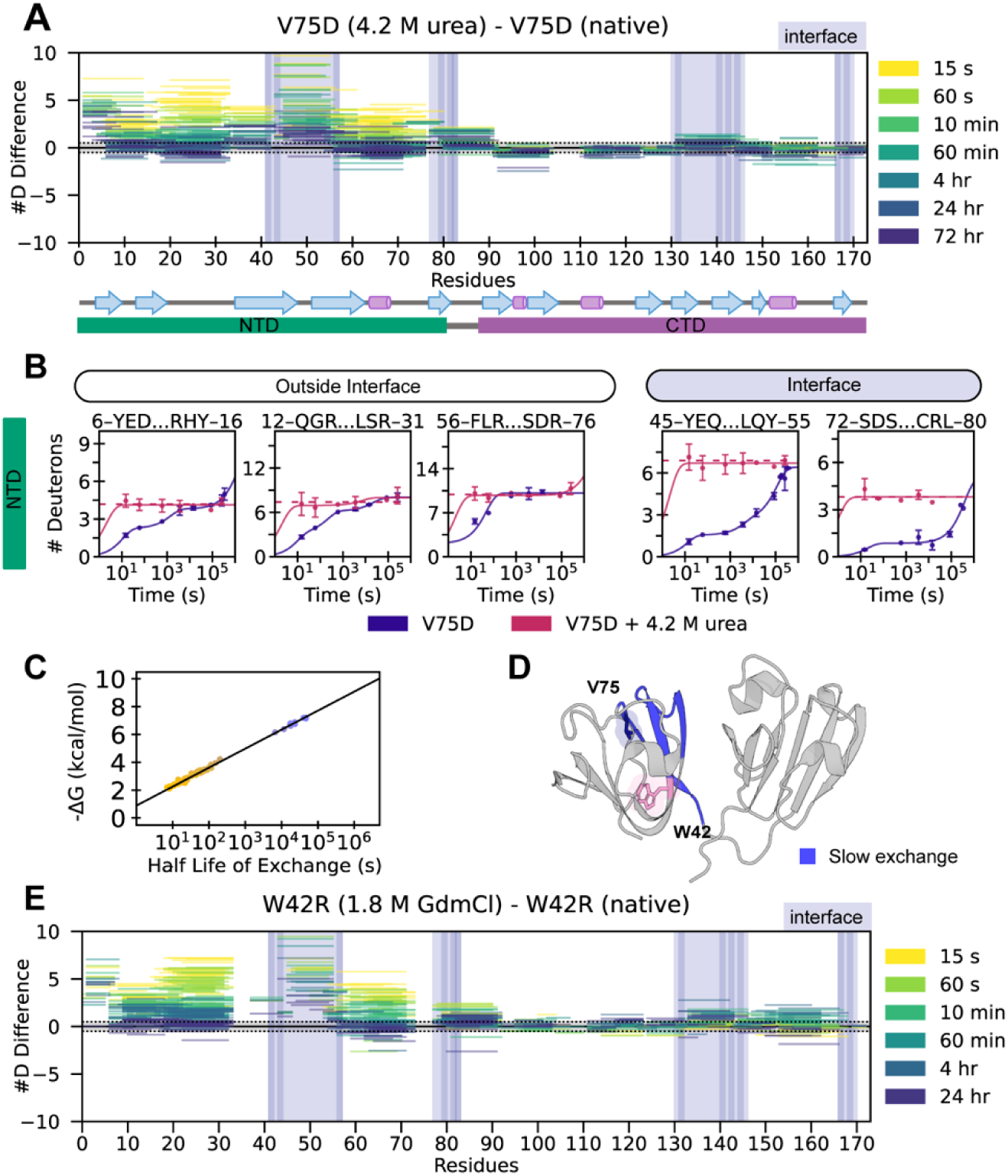
V75D and W42R undergo slow hydrogen-deuterium exchange at the NTD interface. A) Subtractive plot of average deuteration of V75D in urea – deuteration of V75D in native conditions for all timepoints listed. Each line represents an individual peptide spanning the residues indicated on the x-axis, with difference in number of deuterons uptaken indicated on the y-axis. A positive difference in deuteration indicates less deuteration (more protection) under native conditions than in urea. Secondary structure map and domain regions are indicated below the plot. B) Uptake plots for representative peptides in the NTD of V75D, under native conditions (blue) or in 4.2 M urea (magenta). The dotted magenta line is the average of V75D uptake in urea. The solid lines are a multiexponential fit to uptake data. Error bars represent standard deviation of technical replicates. C) Graphical illustration of the relationship between half-life of exchange and estimated ΔG for peptides in EX2. Each point is an individual peptide in the NTD of V75D, where a half-life of exchange is found by fitting uptake data to a multiexponential equation and ΔG_peptide_ is subsequently calculated based on that peptide’s amide intrinsic exchange rates. The black line is the average amide intrinsic exchange rate across the entire NTD, *k_int_* = 6.29 s^-1^. Most peptides (yellow) cluster near or under the ΔG of the NTD unfolding transition of V75D, which is 3.5 ± 0.8 kcal/mol. However, some peptides (blue) exchange at a rate corresponding to a significantly higher ΔG. D) Blue regions highlighted on the crystal structure are regions of unusually slow exchange corresponding to the blue peptides in C. E) Same as A, but depicting data collected from the W42R variant either under native conditions or in 1.8 M GdmCl.

Under native conditions, we identify a different intermediate, a conformation in which some regions of the NTD exchange too slowly to be associated with the exposed-interface intermediate. The expected stability of the exposed-interface intermediate in V75D based on the equilibrium fluorescence studies (3.5 kcal/mol, Table 1) corresponds to an average half-time for exchange of approximately 10 minutes (based on the known intrinsic exchange rates for the amides in the NTD under these conditions (48), Figure 2C). Under native conditions, almost all amides in NTD peptides exchange within this time frame, exhibiting complete exchange at the 60-minute timepoint (Figure 2A-C). However, notably, amides in NTD peptides that contribute to the interface (residues 43-55 and 71-82, Figure 2D) are not completely exchanged after 60 minutes and, in fact, do not exhibit full exchange even after 96 hours (3 x 10^6^ sec), resulting in an estimated half-life of exchange ∼ 10^4^ or 10^5^ seconds. This anomalously slow exchange corresponds to a predicted ΔG value greater than 6 kcal/mol (Figure 2C), inconsistent with expectations for the exposed-interface intermediate. We therefore term the conformation associated with this new intermediate state the “buried-interface intermediate.”

In order to evaluate whether this unusual behavior is unique to V75D, we monitored exchange of another variant with a destabilized NTD, the W42R variant, in which the NTD is destabilized by ∼5 kcal/mol (25). Similar to V75D, the native structure of W42R exhibits no major structural changes from wild type, although a small population of a conformation with partial unfolding in the NTD was observed by solution NMR spectroscopy (25). Using HDX-MS, we find that, like V75D, W42R exhibits unexpected high protection along the interface (Figure 2E), and the exchange behavior of the exposed-interface intermediate (populated at 1.8 M GdmCl) is nearly identical to that of V75D (Figure 2A, E). Thus, W42R’s HD exchange in the NTD is very similar to that of V75D (Figure S4), and both variants populate a buried-interface intermediate under native conditions.

### HDX in the isolated domains

To determine whether the unexpected slow exchange observed along the NTD interface relies on interaction with the CTD, we measured continuous hydrogen exchange behavior in the isolated γDC_NTD_ and γDC_CTD_ domains and compared these to the same regions in the full-length wild-type γDC. For all peptides in γDC_NTD_ and γDC_CTD_ that exhibit EX2 behavior, exchange half-lives were consistent with global unfolding of each domain, including along the regions that comprise the interface in the full-length protein. Amides in peptides along the interface exchange faster in the isolated domains than in full-length γDC (Figure 3A, B). Unfortunately, in the full-length wild-type protein the NTD is too stable to permit monitoring HD exchange on an easily accessible timescale, and therefore no information can be obtained about the putative intermediate interface in this construct.

**Figure 3.**
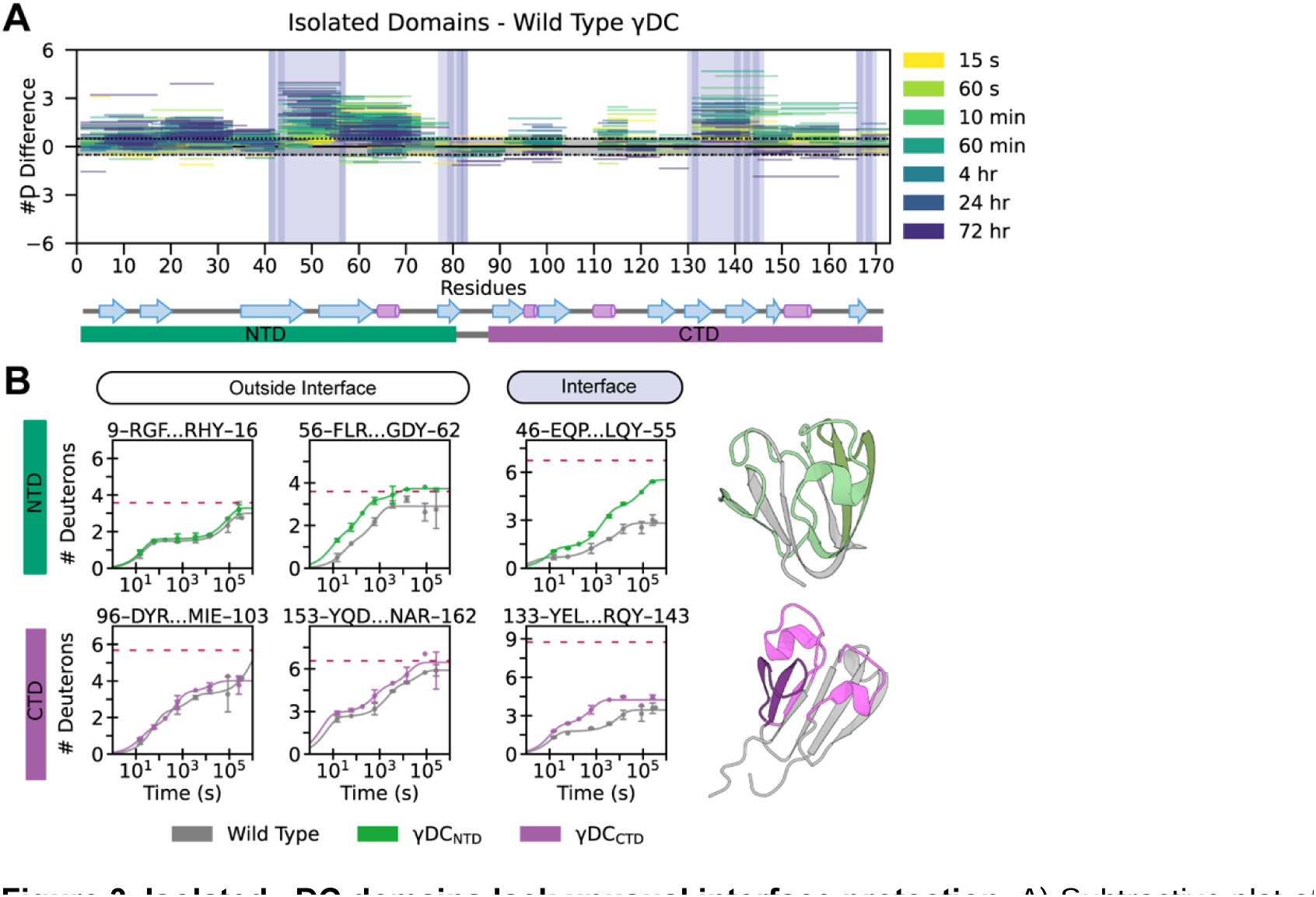
IsolatedγDC domains lack unusual interface protection. A) Subtractive plot of deuteration of each isolated domain (γDC_NTD_ or γDC_CTD_) – wild-type γDC. Highlighted regions on each crystal structure (γDC_NTD_, left; γDC_CTD_, right) indicate notable differences in deuteration. B) Representative uptake plots for comparison of wild-type and isolated domains. The dotted magenta line is average deuteron uptake for V132A in 4.2 M urea (fully exchanged comparison).

### Changes in hydrophobic contact residues in the CTD interface destabilize the buried-interface intermediate

To probe the role of interactions in the CTD interface region with the NTD, we introduced amino acid changes into the CTD and examined the effect on the hydrogen exchange of peptides in the NTD. If interfacial hydrophobic interactions with the CTD are contributing to the stability of the intermediate populated under native conditions, then weakening these interactions by reducing side chain contacts, such as in the V132A variant, should affect the intermediate. We carried out continuous HDX on both the V132A variant and the V75D/V132A variant, to measure the impact of CTD-interface variation in both wild-type and NTD-destabilized contexts.

In the V132A variant, most amides in NTD peptides (with the exception of those at the slow-exchanging interface) exchange at a rate similar to or faster than measured for the same region in the isolated NTD (γDC_NTD_). While exchange in NTD interface peptides is faster for V132A than for wild-type γDC, it is still slower than in the isolated wild-type γDC_NTD_ (Figure 4A, B). This suggests partial, though not complete, loss of interfacial interactions. The V132A CTD exhibits exchange broadly similar to the isolated V132A-γDC_CTD_, except for slightly slowed exchange at the interface. The isolated V132A-γDC_CTD_ exhibits minor loss of protection in peptides near position 132, but otherwise closely resembles the wild-type γDC_CTD_ (Figure S5). Taken together, this suggests that NTD destabilization in the V132A variant is the result of the loss of hydrophobic contacts mediated by the shortening of the hydrophobic side chain at V132, but that weaker stabilizing interactions at the interface persist. This is consistent with our observation from the Ising analysis that in the V132A variant, some stabilization energy at the NTD-CTD interface is retained even when the NTD is unfolded. In an attempt to further weaken the interface, we added four more amino acid changes in the CTD at the interface to yield the dInt-γDC variant (V132A, Q143A, L145A, M147A, and V169A) (30, 37), and we monitored its stability via hydrogen exchange. Exchange rates of amides in peptides from the NTD of dInt-γDC are mostly indistinguishable to those in the NTD of V132A γDC, suggesting that these additional changes do not weaken the interface further and only affect the intrinsic stability of the CTD (Figure S2B). This is further evidenced by the relative stabilities of the two unfolding transitions of dInt-γDC in denaturant (Figure S2C). Therefore, we did not study this variant further, but focused on adding the V75D amino acid change to the V132A variant to explore the role of the interface in the newly identified intermediate accessible under native conditions.

**Figure 4.**
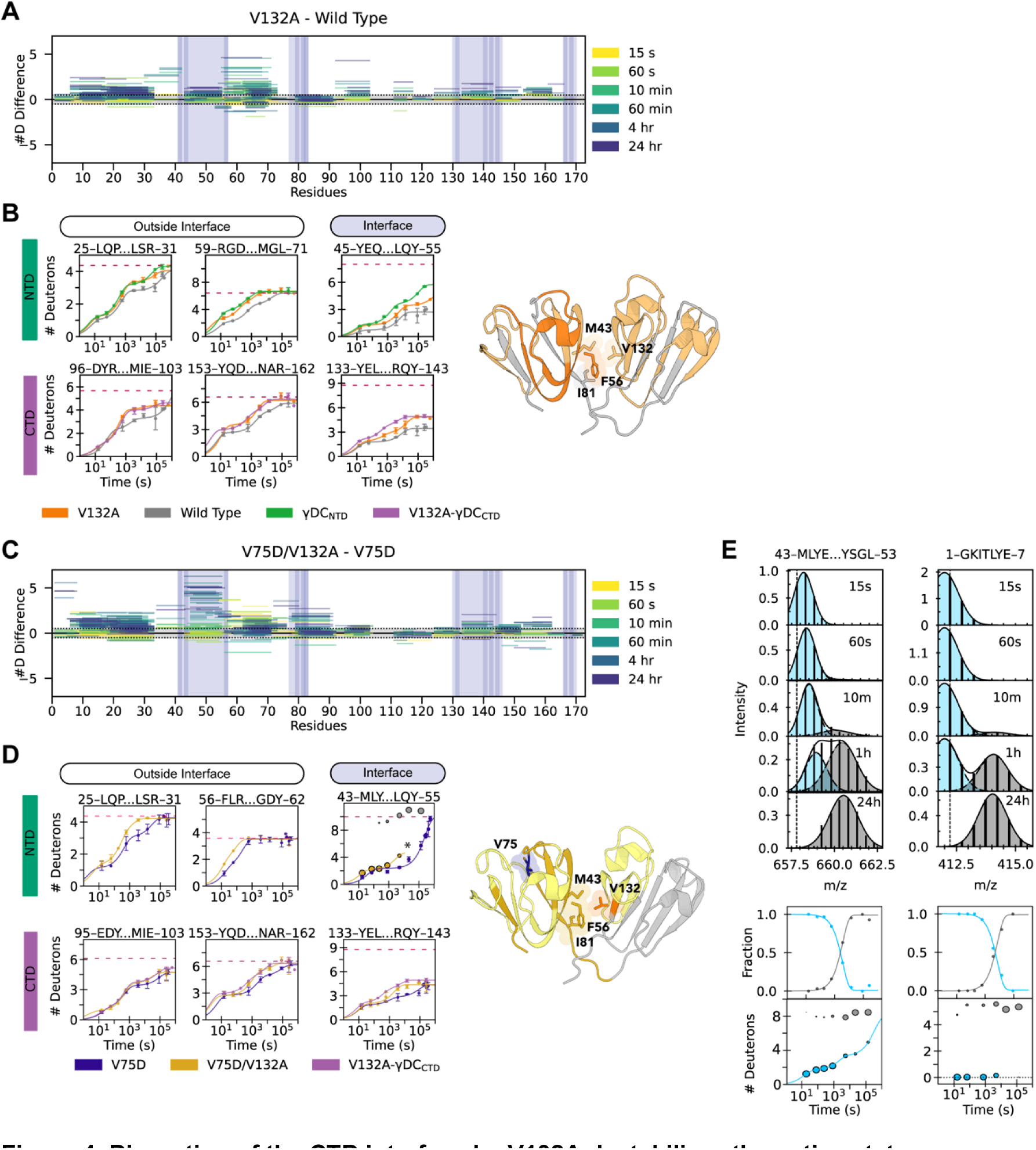
Disruption of the CTD interface by V132A destabilizes the native state intermediate. A) Subtractive plot of deuteration of V132A γDC – wild-type γDC. B) Uptake plots of V132A, wild-type γDC, and isolated domains. The dotted magenta line is average deuteron uptake of V132A in 4.8 M GdmCl (fully exchanged comparison). Highlighted regions on the crystal structure indicate notable differences in deuteration (darker, more difference; gray = no difference). V132A and its three closest interacting residues in the NTD are highlighted. C) Subtractive plot of deuteration of V175D/V132A γDC – V75D γDC. D) Uptake plots depicting V75D (dark blue) and V132A (yellow). The dotted magenta line is V132A in 4.8 M GdmCl (fully exchanged comparison). Peptides at the NTD interface in V75D/V132A demonstrate bimodal behavior and are plotted in yellow and gray to represent the lighter and heavier populations, respectively, where the size of the dot represents the relative population. Highlighted regions on the crystal structure indicate notable differences in deuteration between V75D/V132A and V75D, with dark mustard corresponding to regions that are bimodal in V75D/V132A. E) Top: Representative mass spectra of two peptides in V75D/V132A as time of exchange increases, with overlaid Gaussian fits to indicate two populations (light blue, lighter, less-exchanged population; gray, heavier, more-exchanged population). The first peptide (residues 43-53) is from the interface; the second peptide (residues 1-7) is from the β1 strand. Middle: Kinetics of conversion between the two populations, depicting change in fraction of heavy and light populations over time. Bottom: Deuteration (based on centroid mass) and relative population (depicted as the size of each dot) of the lighter (light blue) and heavier (gray) populations over time.

The V75D/V132A variant exhibits less protection than V75D across the entire NTD, with the most dramatic changes seen at the interface (Figure 4C, D). Notably, amides in peptides at the V75D/V132A interface exhibit bimodal exchange behavior (residues 43-55, 72-80) (Figure 4D, E). This bimodal behavior does not result from only EX1 kinetics (as exemplified for peptide 1-7, Figure 4E), since the centroid (or average mass) of the lighter peak increases with time (peptide 43-53). Therefore, this behavior indicates the presence of two different populations of conformers, one undergoing EX2 exchange and the other undergoing EX1 exchange, with a slow rate of interconversion between the two. The EX2 hydrogen exchange of the lighter population at the interface is slow compared to the rest of the peptides in the NTD, but it is faster than in V75D (Figure 4D). This suggests that some degree of destabilization at the interface is reflected in the EX2 transition, although competition with the EX1 exchange pathway renders an accurate quantification difficult (see below).

### Peptides with EX1 kinetics report on a correlated opening transition

In most experiments, we observe only a few peptides whose amide exchange lies in the EX1 regime. For these peptides, the observed HDX exchange rates report on the kinetics of the opening reaction (a transition from an exchange-incompetent, or closed, state to an exchange-competent, or open, state). Peptides that exhibit this behavior reside in two locations: the first beta strand of the NTD or, under some circumstances, the NTD interface.

The first beta strand of the NTD (residues 1-7, β1) exhibits EX1 behavior in every γDC variant investigated here, including in the isolated γDC_NTD_, indicating the presence of slow interconversion between the open and closed states involving this strand (Figure S6). Interestingly, neither of β1’s hydrogen-bonding partners, β2 and β4 in the NTD β sheet, shares this EX1 behavior. The measured opening rate of the β1 strand increases upon destabilization of the NTD, whether by amino acid substitution or by addition of denaturant (Figure 5A, 5B, Table 2). When measured in V75D against urea concentration (0.6 M - 3.0 M urea), the natural logarithm of this rate increases linearly (Figure 5C).

**Figure 5.**
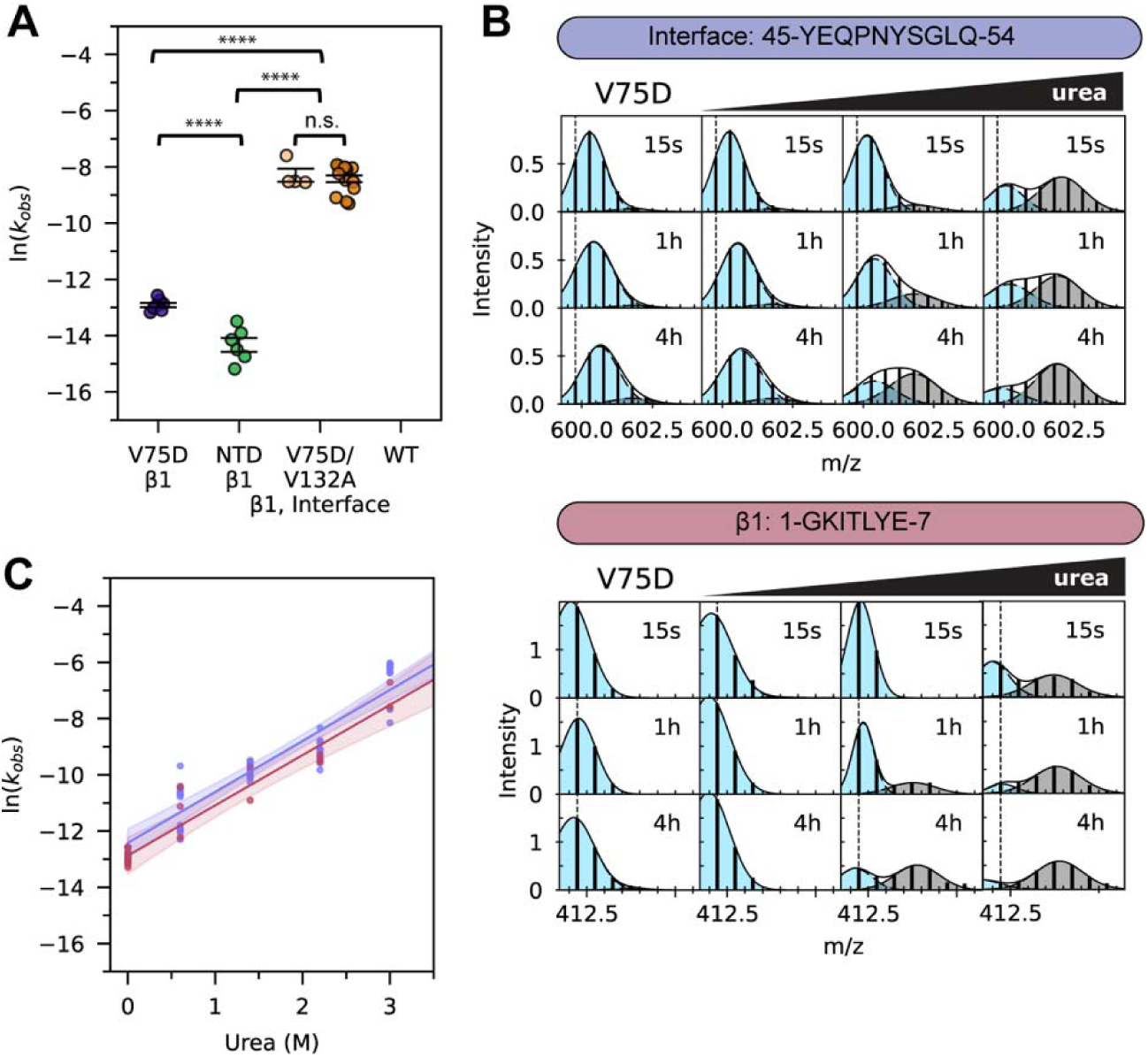
Rates of exchange at β1 and the buried interface correlate. A) Comparison of rates of EX1 β1 exchange in V75D (dark blue), NTD (green), and V75D/V132A (light orange), along with EX1 interface exchange in V75D/V132A (dark orange). Error bars indicate standard error of the mean. Asterisks indicate *p* < 0.0001 by ANOVA and Tukey HSD. B) Example mass spectra of V75D peptides in β1 and at the interface when equilibrated in increasing amounts of urea. C) Rates of V75D at both β1 (light red) and the NTD interface (slate blue). Shading represents 95% CI. The linear increase in EX1 opening rates of the V75D interface with respect to urea is indistinguishable from the linear increase in rates of opening at V75D β1 (p = 1.0 by one-way ANCOVA).

**Table 2.** Apparent EX1 HDX rates of exchange. Rates (s^-1^) were measured at two different regions in γDC variants, the β1 strand (residues 1-7) and the interface (residues 43-55), reported as the mean of the natural log of rates (s^-1^) obtained from distinct peptide population transitions individually fit to exponential kinetics. Error reported is standard error of the mean. Peptides were chosen from two (V75D/V132A, γDC_NTD_) or four (V75D γDC) independent exchange experiments. ^(a)^ Wild-type and V132A γDC exchanged too slowly to observe any transition in EX1 on the timescale of our experiments.

| | | $\ln(k_{op})$ | # peptides |
| --- | --- | --- | --- |
| β1 strand | **V75D γDC** | $-12.9 \pm 0.08$ | 7 |
| | **γDC<sub>NTD</sub>** | $-14.3 \pm 0.25$ | 6 |
| | **V75D/V132A γDC** | $-8.3 \pm 0.23$ | 4 |
| | **Wild-type γDC** | $< -15$ | (a) |
| | **V132A γDC** | $< -15$ | (a) |
| Interface | **V75D/V132A γDC** | $-8.4 \pm 0.12$ | 15 |

We also investigated the exchange behavior of a small peptide that comprises the amino acid sequence of the first 10 amino acids of γDC. Exchange is rapid for this peptide in isolation (complete exchange < 15 s), indicating that this peptide does not exhibit any intrinsic structure. Therefore, we posit that the behavior of β1 reports on the opening kinetics of the exposed-interface intermediate. The EX1 behavior is consistent with the kinetics of folding/unfolding of the isolated γDC_NTD_ (33). Under our experimental conditions, using a two-state model, extrapolation of denaturant unfolding yields an approximate folding rate of ∼ 0.47 s^-1^ (Figure S7). Given the limitations of such extrapolations, this rate is within an order of magnitude slower than the intrinsic rate of exchange (average k_int_ for this peptide ∼ 3.2 s^-1^). This extrapolated unfolding rate is faster than rates of exchange as measured via hydrogen exchange at β1 (Table 2). This is to be expected if unfolding intermediates are present (49), as has been previously suggested (26, 31).

Interestingly, the measured EX1 rates of amides in β1 are equivalent to rates observed for the EX1 component in the bimodal exchange at the NTD interface in the V75D and V75D/V132A variants. A combination of EX2 and EX1 exchange at the V75D NTD interface is observed when the interface is destabilized, either by amino acid substitution (as previously noted in V75D/V132A, Figure 4E) or by addition of urea (Figure 5B). Under both circumstances, rates of EX1 opening in peptides at the NTD interface correlate strongly with the measured rates of opening at β1. The opening rate for amides of the V75D/V132A variant at the NTD interface (ln(*k*) = -8.4 ± 0.12) is indistinguishable from the opening rate for β1 (ln(k) = -8.3 ± 0.23) (Figure 5A, Table 2), and in urea, opening rates at the V75D NTD interface are indistinguishable from opening rates of V75D β1 (p = 1.0 by one-way ANCOVA) (Figure 5C). EX1 exchange in NTD interface amides is possibly occurring in variants other than V75D/V132A; however, in most other constructs, observation of EX1 exchange in peptides at the interface in the exposed-interface intermediate is prohibited by the interfering observation of slow EX2 exchange in the buried-interface intermediate.

The equivalence of EX1 exchange rates observed for amides in β1 peptides and in peptides at the NTD interface demonstrates that amides in β1 and at the interface are likely reporting on the same transition. Therefore, we conclude that the EX2 exchange at the interface is associated with the buried-interface intermediate, while the faster-exchanging EX1 population is associated with the transition to the exposed-interface intermediate (Figure 4E, 5B).

## Discussion

Many proteins associated with aggregation-related diseases have a common feature: the ability to populate a non-native or partially unfolded conformation that is more aggregation-prone than the native state (36). Characterizing these conformations and the means by which environmental and sequence factors influence this mis- or partial folding is therefore critical to mechanistically explain these diseases. Using a combination of hydrogen-deuterium exchange/mass spectrometry and traditional chemical denaturation, we have identified and characterized intermediates on the energy landscape of γD-crystallin under both native and denaturing conditions. We find that, for two cataract-prone variants of γD-crystallin, the primary intermediate populated under native conditions is structurally and energetically distinct from the intermediate populated under chemically denaturing conditions. This intermediate involves partial unfolding of the NTD, as compared to the full unfolding which is known to occur under mildly denaturing conditions, and it is therefore hidden to structural and biochemical analyses that require perturbation using denaturants. This newly described intermediate, the buried-interface intermediate, consists of a partially folded NTD and a fully folded CTD.

Our model for the buried-interface intermediate is based on the unexpected hydrogen exchange behavior we observe in the regions of the NTD that form the interface with the CTD. Given the thermodynamic importance of the NTD/CTD interface to NTD stability, especially in variants with an otherwise highly destabilized NTD, we expected that these interface peptides would be among the most protected regions of the NTD, but that they would either exchange at a rate consistent with the stability of the NTD unfolding transition, or else exchange in the EX1 regime if the NTD interface is slow to refold. Instead, we found that peptides along the interface in the NTD are too highly protected in EX2 for their opening transition to correspond to full unfolding of the NTD, implying the presence of an alternative intermediate population containing some degree of structure at the interface. We observe this buried-interface intermediate in both NTD-destabilized variants studied, V75D and W42R.

We do not observe anomalously slow EX2 exchange in the isolated N-terminal domain fragment γDC_NTD_. This demonstrates that the unusually slow interface exchange only takes place in full-length γDC, and that the slow exchange behavior along the NTD interface is a consequence of interaction with the CTD, rather than being a consequence of some native misfolding or interaction event taking place within the NTD itself.

The interface of γDC is critical to its stability and unfolding rate and is crucial for the folding and stability of the NTD – particularly in NTD-destabilized variants. In addition to quantifying the strength of this interface by an Ising analysis, we here demonstrate that residues at the interface are involved in a buried-interface equilibrium intermediate. Moreover, we find that the population of this intermediate, compared to the exposed-interface intermediate, can be modulated by addition of denaturant or by amino acid changes at the CTD side of the interface, as seen in V132A variants. Disrupting the interface enables direct observation of both intermediates simultaneously. Surprisingly, we also prove that interfacial interactions in the V132A variant, though weaker than in wild-type, persist even when the NTD is unfolded.

Our results permit us to propose a model for V75D γDC’s energy landscape (Figure 6). The buried-interface intermediate (BII) has both a free energy closer to that of the native state and a lower kinetic barrier than the exposed-interface intermediate (EII) present under denaturing conditions. The observation of EX1 transitions in β1 and at the interface suggests two possible models: one in which BII is not on the unfolding pathway to EII (EII ⇌ N ⇌ BII, Model 1) (Figure 6A), and one in which it is (N ⇌ BII ⇌ EII, Model 2) (Figure 6B). We favor Model 1 as the most likely scenario, as outlined below.

**Figure 6.**
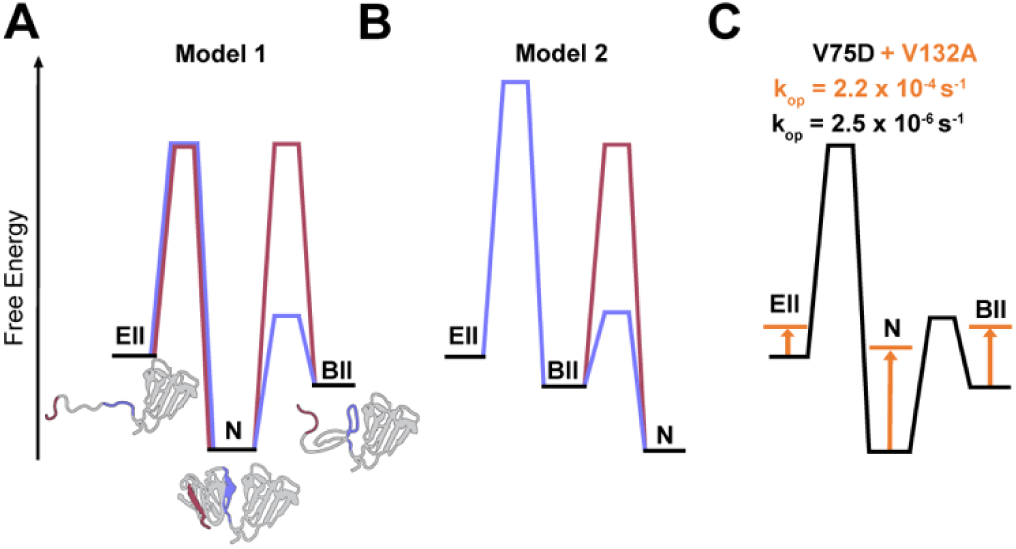
The energy landscape of V75DγDC. A, B) Two potential placements of the buried-interface intermediate, on-pathway (Model 1) and off-pathway (Model 2). The colored kinetic barriers (large or small) correspond to the observation of EX1 (large) or EX2 (small) exchange at either the interface (slate blue) or β1 (light red). In Model 1, unfolding occurs through two sequential barriers which must be equivalent, with BII on-pathway. In Model 2, unfolding occurs through either the N ➔ EII or N ➔ BII pathway. C) In a moderately denaturing context, NTD-destabilized γDC (V75D or W42R) populates the exposed-interface intermediate. Under native conditions, the most-populated intermediate is the buried-interface intermediate, with a high kinetic barrier restricting access to the exposed-interface intermediate. V132A’s impacts on the energy landscape are illustrated in orange (some effects are estimated). Destabilization of the interface either by amino acid substitution or by mild denaturant lowers the kinetic barrier to population of the exposed-interface intermediate, permitting observation of both intermediates at once.

β1 is in EX1 exchange under all conditions, which implies a high barrier to β1 closing in both BII and EII. However, peptides at the interface only undergo EX1 exchange when unfolding to the EII. In Model 2, there are two sequential barriers, both of which show EX1 exchange at β1. The closing rates of β1 for both barriers (EII ➔ BII and BII ➔ N) must be slow, or else we would observe EX2 exchange at β1 in one of those states. However, for peptides at the interface, only the EII ➔ BII closing rate is slow, because EX2 behavior is observed in BII. This implies that EX1 exchange at interface peptides specifically report on the rate of opening BII ➔ EII. Since EX1 exchange rates at β1 correlate with rates of EX1 exchange at the interface, the BII ➔ EII and the N ➔ BII opening rates are indistinguishable in this model. In Model 1, on the other hand, the two barriers (N ⇌ BII and N ⇌ EII) are not sequential, consistent with EX1 exchange N ➔ EII at both β1 and the interface. For the N ➔ BII transition, the interface remains protected and subsequently exchanges in EX2 to some other open state. β1 unfolds but exhibits a slow closing rate resulting in EX1 exchange. The relative population of each intermediate can be altered by the addition of denaturant or by amino acid substitution at the interface (Figure 6C). This results in a lower kinetic barrier to interface unfolding, permitting the simultaneous observation of both intermediates.

While this work cannot determine whether the identified hidden intermediate is aggregation-prone, or whether a direct link between interface stability and aggregation exists, it is highly suggestive that the buried-interface intermediate plays a role along the pathway to aggregation and cataract formation. Such equilibrium intermediates have been shown to be crucial precursors in aggregation pathways for various diseases (50). Moreover, partially folded species may be relevant conformers encountered and bound by α-crystallins (51). While *in vitro* aggregation pathways have been previously described for γDC upon rapid refolding from high concentrations of denaturant (28), such conditions are not encountered *in vivo*. In the context of aggregation from native conditions, it is important to consider excursions from the native side of the folding barrier, particularly when the kinetic barriers to unfolded states are very high.

The observation of differential stability between γDC’s two domains has led to a suggestion of domain swapping as a possible mechanism for γDC aggregation (29). Contrary to a simple domain-swapping model, in which the entire NTD and CTD may be exchanged between monomers, the partially unfolded intermediate uncovered and characterized here retains substantial structure at the NTD-CTD interface. The increased mobility and conformational variability across the NTD caused by a destabilizing amino acid change, combined with a local stabilizing effect of the NTD-CTD interface, results in a partially unfolded NTD, which exposes a surface that normally is buried in the full-length structure and in folded γDC_NTD_. Thus, we suggest that it is essential to search for and describe equilibrium intermediates under experimental conditions that emulate physiological circumstances.

## Materials and Methods

Detailed procedures are provided in SI Appendix.

### Expression and purification of γD-crystallin variants

Mutations in the gene for wild-type γD-crystallin (from (52)) were created via site-directed mutagenesis (IDT). Mutations were confirmed by Sanger sequencing (Quintara). Wild-type and all variant γD-crystallins except for the V75D/V132A double variant were purified as described previously (32).

### Determination of global stability by intrinsic tryptophan fluorescence

For each construct, two 5 μM protein stocks were prepared: a no denaturant protein stock and a high GdmCl (5.6 M) protein stock, both in PBS pH 7.0 (Sigma-Aldrich P4417), 5 mM DTT. Samples with a range of GdmCl concentrations were prepared by combination of the two stocks and allowed to equilibrate at room temperature for at least 24 hours for all constructs except wild-type, which was equilibrated for 172 hours. Measurements were then performed at 25 °C using a PTI Quantamaster Fluorometer (Horiba). An excitation wavelength of 280 nm was used to excite tryptophan residues, and emission spectra were recorded from 310 to 390 nm (0.7 s/nm).

Samples were recovered from the cuvette after each measurement and the exact GdmCl concentration was determined by taking the refractive index (53). Signal was reported as a ratio of signal at 360 nm to signal at 320 nm. Signal ratios per concentration GdmCl from each variant were fit to a two-state or three-state folding model using Python’s LMFIT module (54), which allowed determination of the transition midpoint, ΔG_unfolding_ and *m*-value for each transition.

To globally fit unfolding transitions of γDC variants, we generated fitting equations using a one-dimensional Ising model as described previously (39, 55). Python code modified from Marold *et. al.* (55) was used to generate and fit partition functions describing the fraction of a given folded state as a function of denaturant using separate equilibrium constants for folding of individual domains and for the interface coupling (Table S3). To fit V132A data, we modified the 1-D Ising partition function to permit coupling between the CTD and the unfolded NTD.

### HDX-MS continuous exchange

Deuterated buffers were prepared by lyophilizing PBS pH 7.0 containing 5 mM DTT and resuspending in D_2_O (Sigma-Aldrich 151882). All urea- and GdmCl-containing buffers were lyophilized and deuterated a total of three times to ensure total deuteration of the denaturant. Protein samples to be exchanged in the presence of denaturant were equilibrated by incubation in the requisite concentration of denaturant at 25 °C in PBS pH 7.0, 5 mM DTT for at least 24 hours. The peptide angiotensin-II (sequence DRVYIHPF, Thermo Scientific) was included in all samples at a concentration of 0.25 μg/mL as a fiduciary to correct for potential variability in back exchange between different buffers.

To initiate continuous labeling, samples were diluted tenfold into temperature-equilibrated, deuterated PBS buffer to produce a final γDC concentration of 7.5 μM. Samples were quenched at 15 s, 60 s, 10 min, 1 hr, 4 hr, 24 hr, and 72 hr by mixing 6 μL of the partially exchanged protein with 24 μL of quench buffer 1 (8.6 M urea, 500 mM TCEP pH 2.2) on ice. Extra time points were collected in some experiments for additional resolution. Quenching samples were incubated on ice for 1 minute to allow for partial unfolding to assist with proteolytic degradation and then were flash frozen in liquid nitrogen and stored at −80D°C. Samples were thawed by resuspension with 50 μL of ice-cold quench buffer 2 (0.75 M glycine, 50 mM TCEP, pH 2.2) to reduce denaturant concentration prior to proteolytic digestion and immediate injection onto the liquid chromatography-mass spectrometry system. Inline digestion was performed with aspergillopepsin (Sigma-Aldrich P2143) and porcine pepsin (Sigma-Aldrich P6887).

### HDX-MS data analysis

Briefly, γDC peptides were identified using Byonic software (Protein Metrics) and subsequently, isotopic distributions of each peptide were fit and verified manually using HDExaminer3 (Sierra Analytics). For unimodal isotopic distributions, the monoisotopic mass for each peptide was subtracted from the mass centroid and extracted. Bimodal isotopic distributions were exported from HDExaminer3 and globally fit to a sum of two Gaussian distributions. All downstream quantitative analysis was performed using Python scripts in Jupyter notebooks.

## Author Contributions

S.V., A.J.G, A.M.G, and S.M. designed research; S.V carried out experimental studies with assistance from J.R.M.; S.V. and S.M. analyzed data; S.V., A.J.G, A.M.G, and S.M. wrote and edited the manuscript.

## Acknowledgments

We thank the entire Marqusee lab for experimental advice and support. This work was supported by funding from the NIH (SM), the NSF (SM), and the NEI (AMG). SM is a Chan-Zuckerberg Biohub investigator.

## Notes

### Competing Interest Statement

The authors have declared no competing interest.

