## Supplemental Information for "Cataract-prone variants of γD-crystallin populate a conformation with a partially unfolded N-terminal domain under native conditions"

**This PDF file includes:**

Supporting text
Figures S1 to S7
Tables S1 to S3
SI References

**Supporting Information Text**

**Methods**

*Expression and purification of γD-crystallin variants*

For each construct, an *Escherichia coli* BL21 (DE3) colony bearing a pET14b vector encoding the protein of interest was used to inoculate 50 mL of Luria-Bertani (LB) medium supplemented with 100 mg/mL ampicillin. Following overnight growth at 37˚C in a rotary shaker, this culture was used to inoculate 2 L of fresh LB medium at 37˚C in a rotary shaker to an OD600 of ~0.6 and induced with 1 mM isopropyl b-D-1-thiogalactopyranoside for 3 h at 37˚C. Cells were pelleted and resuspended in ~30 mL of buffer Q (50 mM Tris pH 8.0, 1 mM EDTA, 1 mM dithiothreitol (DTT)) supplemented with 1× Halt protease inhibitor cocktail (Thermo) and benzonase (Novagen), and stored at 80 ˚C prior to thawing for protein purification.

Thawed pellets were lysed by sonication, and the lysate was clarified by centrifugation at 15,000 x g for 30 min. The supernatant was filtered with a 0.2 um vacuum filter and passed over a 5 mL HiTrap Q HP anion exchange column (GE Healthcare Life Sciences, Piscataway, NJ) equilibrated in buffer Q. γDC was collected in the flow-through and diluted 1:1 with buffer S (25 mM MES pH 6.0, 1 mM EDTA, 1 mM DTT, and 2% v/v glycerol), after which the pH was adjusted to 6.0 via dropwise addition of HCl. This eluate was filtered with a 0.2 um vacuum filter and loaded onto a HiPrep SP XL 16/10 cation exchange column (Cytiva) equilibrated in buffer S. Bound proteins were eluted using a 0–75% gradient of 1 M NaCl in buffer S. The crystallin-containing elution peak (as assessed by SDS-PAGE) was passed over either a S75 16/60 size-exclusion column (GE) or S75i 10/300 size-exclusion column (GE) pre-equilibrated in PBS pH 7.0, 5 mM DTT for final purification.

To avoid co-purification of truncated V75D/V132A γDC expression products lacking the NTD, $His_6$-MBP with a C-terminal tobacco etch virus (TEV) proteolysis site fused to V75D/V132A γD-crystallin (pSV040) was cloned via Gibson assembly and grown and expressed as above. Clarified lysate was allowed to batch bind to HisPur $Ni^{2+}$-NTA resin (Thermo) washed with 50 mM Tris pH 7.5, 150 mM NaCl, 20 mM imidazole, 0.5 mM TCEP, and was eluted with 50 mM Tris pH 7.5, 150 mM NaCl, 500 mM imidazole and 0.5 mM TCEP. Eluate was concentrated in an Amicon spin concentrator (Millipore) and loaded onto a Superdex200 16/60 size-exclusion column (GE) equilibrated in 50 mM Tris pH 7.5, 150 mM NaCl, 0.5 mM TCEP. Peaks corresponding to the MBP-crystallin fusion were collected and digested overnight at 4 ˚C with TEV protease with an N-terminal 6× histidine tag, yielding a final product with no scar. The $His_6$-tagged TEV protease and the $His_6$-MBP scaffold were removed via a subtractive $Ni^{2+}$-NTA affinity step using HisPur $Ni^{2+}$-NTA resin pre-equilibrated with 50 mM Tris pH 7.5, 150 mM NaCl, 20 mM imidazole, 0.5 mM TCEP. Flow through was then concentrated and loaded onto an S75i 10/300 size-exclusion column (GE) pre-equilibrated in PBS pH 7.0, 5 mM DTT. Peaks corresponding to the correct crystallin constructs were verified by MS, quantified by UV-Vis absorption at 280 nm, and flash-frozen for storage at -80 ˚C for future use.

*Fitting intrinsic tryptophan fluorescence*

Two-state fit (1):

$$y_{obs} = \frac{F_{intercept} + F_{slope}x + \left(U_{slope} + U_{intercept}x\right)e^{m(x-C_m)/(RT)}}{1 + e^{m(x-C_m)/(RT)}} \quad (1)$$

Parameter definitions are as follows: $F_{intercept}$, folded intercept; $F_{slope}$, folded baseline slope; $U_{intercept}$, unfolded intercept; $U_{slope}$, unfolded baseline slope; $m$, $m$-value; $C_m$, transition midpoint. R is the gas constant and T is temperature. Baseline slopes were permitted to vary.
Three-state fit (2):

$$y_{obs} = \frac{F_{intercept} + F_{slope}x + \left(I_{slope} + I_{intercept}x\right)e^{m_1(x-C_{m1})/(RT)} + \left(U_{slope} + U_{intercept}x\right)e^{m_1(x-C_{m1})/(RT)}e^{m_2(x-C_{m2})/(RT)}}{1 + e^{m_1(x-C_{m1})/(RT)} + e^{m_1(x-C_{m1})/(RT)}e^{m_2(x-C_{m2})/(RT)}} \quad (2)$$

Parameter definitions are as follows: $F_{intercept}$, folded intercept; $F_{slope}$, folded baseline slope; $I_{intercept}$, intermediate unfolded intercept; $I_{slope}$, intermediate baseline slope (held at zero for all constructs but V132A); $U_{intercept}$, unfolded intercept; $U_{slope}$, unfolded baseline slope; $m_1$, transition one *m*-value; $C_{m1}$, transition one midpoint; $m_2$, transition two *m*-value; $C_{m2}$, transition two midpoint. R is the gas constant and T is temperature.

The 1-D Ising partition function used to fit wild-type data contains terms for four states: unfolded, only NTD folded, only CTD folded, and both NTD and CTD folded and interacting (Table S3**Error! Reference source not found.**). We incorporate native and denatured baselines (locally fit) by multiplying baselines by fraction-folded expressions. To fit V132A data, we modified the 1-D Ising partition function to permit coupling between the CTD and the unfolded NTD. It still contains terms for four states—unfolded, only NTD folded, only CTD folded and interacting with the unfolded NTD, both NTD and CTD folded and interacting—but an additional equilibrium constant is included to describe coupling between the unfolded NTD and the folded CTD: $\omega_C = e^{-\Delta G_{unfoldedN,C}/(RT)}$, which we held equivalent to $\tau_{N,C}$ for this construct. In order to include an intermediate baseline term, we separated the fraction-folded expression into three terms which are multiplied by their respective baseline parameters (locally fit) (Table S3).

*LC-MS*

HDX samples thawed by resuspension were immediately injected into a cooled valve system (Trajan LEAP valve box) connected to a Thermo Ultimate 3000 LC and Q-Exactive Orbitrap MS using a 250 µL sample loop. Inline digestion was performed with firstly aspergillopepsin (Sigma-Aldrich P2143) and secondly porcine pepsin (Sigma-Aldrich P6887) conjugated to beads (Thermo Scientific POROS 20 Al aldehyde activated resin 1602906) and packed into homemade protease columns. After digestion, peptides were desalted over a hand-packed trap column (Thermo Scientific POROS R2 reversed-phase resin 1112906, 1 mm ID × 2 cm, IDEX C-128). Digestion and desalting took place over 6 minutes: 4 minutes at a flow rate of 50 µL/min, and 2 minutes at a flow rate of 300 µL/min.

Acetonitrile, formic acid, and MS-grade water (Fisher Optima LC/MS) were used to prepare mobile phase solvents A (0.1% formic acid) and B (100% acetonitrile, 0.1% formic acid). Peptides were separated by a linear gradient from 5–40% solvent B over a C8 analytical column (Thermo Scientific BioBasic-8 5-µm particle size 0.5 mm ID × 50 mm 72205-050565) with a guard column attached to the inlet (Thermo Fisher Scientific) for 12 minutes, followed by 40–90% solvent B over 30 seconds. Following peptide elution, analytical and trap columns were subjected to a sawtooth wash and subsequently equilibrated at 5% solvent B prior to the next injection. Protease columns were washed with two (for deuterated samples) or three (for undeuterated samples) injections of 100 µL 1.6 M GdmCl, 0.1% formic acid (passed through a 0.2µm filter). Peptides were eluted directly into a Q Exactive Orbitrap Mass Spectrometer operating in positive mode (resolution 140,000, AGC target $3 \times 10^6$, maximum IT 200 ms, scan range 300–1,500 *m/z*). A tandem mass spectrometry experiment was performed for undeuterated samples of each crystallin construct on every day that data for that construct was collected, in order to generate corresponding peptide lists and retention times (full MS settings the same as above, dd-$MS^2$ settings as follows: resolution 17,500, AGC target $2 \times 10^5$, maximum IT 100 ms, loop count 10, isolation window 2.0 *m/z*, NCE 28, charge state 1 and ≥7 excluded, dynamic exclusion of 15 seconds). LC and MS methods were run using Xcalibur 4.1 (Thermo Scientific).

*HDX-MS data analysis*

Peptides were identified using Byonic software (Protein Metrics) with the γD-crystallin sequence containing the corresponding mutations and the sequence of angiotensin-II used as the search library. Sample digestion parameters were set to non-specific; precursor mass tolerance and fragment mass tolerance were set to 6 and 40 ppm, respectively. Peptide lists were imported into HDExaminer3 (Sierra Analytics) along with the deuterated and undeuterated sample mass spectra for analysis. Isotopic distributions were fit and checked manually using HDExaminer3; the monoisotopic mass for each peptide was subtracted from the mass centroid and extracted. Bimodal isotopic distributions were exported from HDExaminer3 for further analysis. Downstream quantitative analysis was performed using Python scripts in Jupyter notebooks.
Bimodal peptide mass spectra for all timepoints were globally fit to a sum of two Gaussian distributions where the heavier peak corresponds to the open state and the lighter peak corresponds to the closed state. The centroid and width of each Gaussian was allowed to vary by only 0.5 Da in cases where EX1 behavior was seen. The center and width of the Gaussians were allowed to vary as needed in cases where a distribution centroid was seen to be migrating. Fractional populations at each timepoint were calculated as the area under each Gaussian and fit to a single-exponential kinetics (equation 3) using the LMFIT package to obtain rates of opening (3).

$$y = h - A\mathrm{e}^{(-kt)} \tag{3}$$

Parameter definitions are as follows: A, amplitude; h, height; $k$, rate ($s^{-1}$).
Uptake data in the EX2 regime were corrected for conditions between different buffers by normalization to exchange of a fiduciary peptide included in all exchange experiments, angiotensin-II (equation 4) (4).

$$D_{corr}(t) = \frac{m(t) - m_{0\%}}{A(t) - A_{0\%}} \tag{4}$$

Parameter definitions are as follows: $m(t)$, peptide centroid mass at a given time; $m_{0\%}$, non-deuterated peptide centroid mass; $A(t)$, angiotensin-II centroid mass at a given time (maximally labeled before 15 seconds); $A_{0\%}$, non-deuterated angiotensin-II centroid mass.
Corrected data were then fit to a multiexponential model (equation 5) using LMFIT (3). For each peptide, either one, two, or three multiexponential terms were used (terms for A, A and B, or A, B and C) (5); inclusion of additional terms in the multiexponential fit was automatically judged on a per peptide basis based on comparison of the Akaike information criteria (AIC) for each model. Each additional term was only included if it decreased the AIC.

$$D(t) = maxP - Ae^{(-k_A t)} - Be^{(-k_B t)} - Ce^{(-k_C t)} - NE \tag{5}$$

Parameter definitions are as follows: D(t), deuteration; maxP, maximum theoretical exchangeable protons (length of the peptide – 2 – number of prolines); A, B, C: number of fast-, medium-, and slow-exchanging protons; $k_A$, $k_B$, $k_C$: rates of exchange for fast-, medium-, and slow- exchanging protons; NE: number of non-exchanging protons.
This permitted calculation of half-lives by finding the fit's intersection at half-maximum exchange, as determined by a corresponding fully exchanged control (or by comparison to the fit's calculated maximum exchange). Half-lives found in this manner were used to estimate expected ΔG values for comparison to states identified by equilibrium denaturation melts by using intrinsic rates calculated using *SPHERE* at www.fccc.edu/research/labs/roder (6).

*Unfolding kinetics of γ$DC_{NTD}$*

A 100 µM stock of γ$DC_{NTD}$ was prepared in PBS pH 7.0, 5 mM DTT, 25 °C. Protein was diluted tenfold in PBS pH 7.0, 5 mM DTT containing varying concentrations of GdmCl (3 M to 5.2 M; the exact GdmCl concentration was determined by taking the refractive index). Dead time between mixing and recording data was 15 seconds. Measurements were performed at 25 °C using a Fluoromax-3 (Horiba). An excitation wavelength of 280 nm was used to excite tryptophan residues, and emission was collected at 350 nm (1 s integration time) until a maximum was reached (~5-20 minutes). Unfolding experiments at each concentration of denaturant were repeated three times. Signal as a fraction of maximum intensity was fit to a single-exponential equation (equation 3) to obtain rates of unfolding.

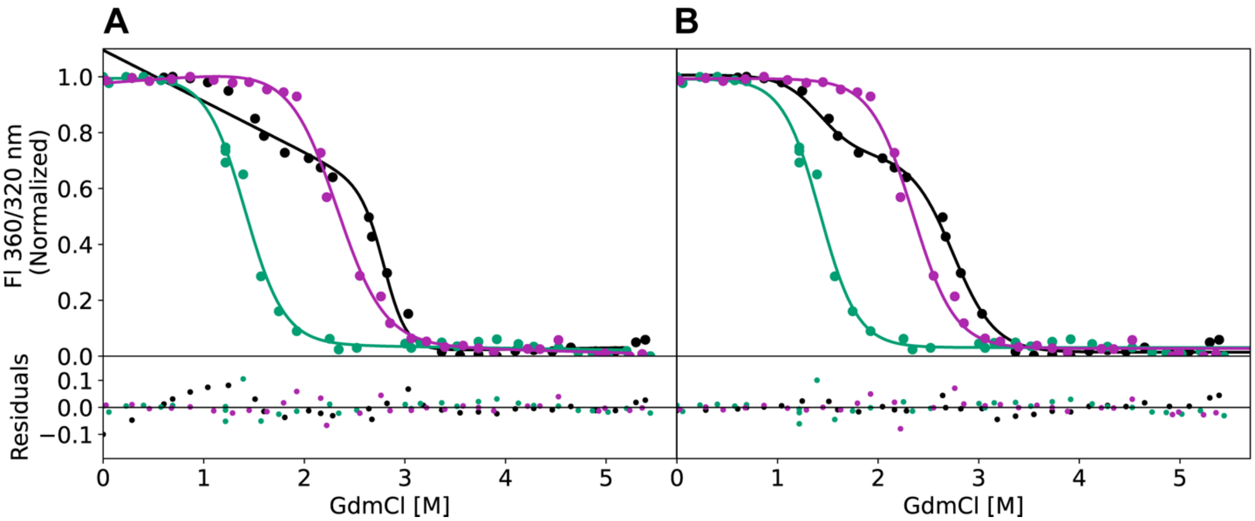


**Figure S1: Comparison of 1D- and modified 1D-Ising models for V132A γDC.** A) V132A γDC equilibrium denaturation data fitted with a 1D-Ising model (γ$DC_{NTD}$, green; V132A-γ$DC_{CTD}$, purple; V132A γDC, black). B) V132A γDC equilibrium denaturation data fitted with a 1D-Ising model modified to include a stabilizing interaction between the unfolded NTD and the folded CTD.

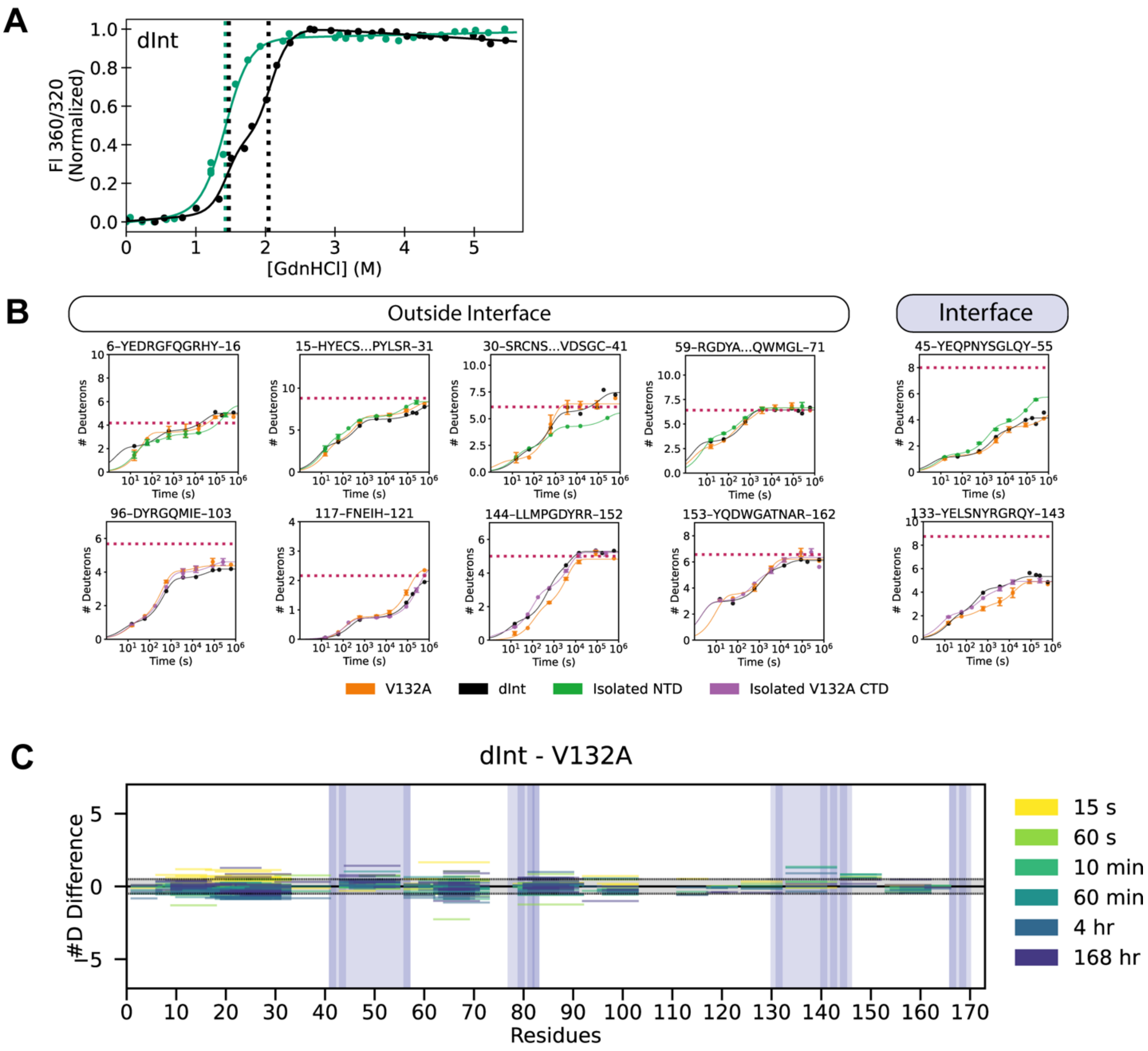


**Figure S2: dInt-γDC amino acid changes destabilize the CTD transition without further impacting the NTD relative to V132A.** A) Equilibrium denaturation of dInt-γDC (green, γDC$_{NTD}$; black, dInt-γDC). B) Uptake plots comparing dInt-γDC, V132A γDC, γDC$_{NTD}$, and V132A γDC$_{CTD}$. The dotted magenta line is average deuteron uptake of V132A in 4.8 M GdmCl (fully exchanged comparison). Exchange of peptides in the NTD of dInt-γDC is mostly indistinguishable to V132A alone: as in V132A, most of the NTD in this full-length construct resembles the isolated γDC$_{NTD}$, while exchange at the interface is more rapid than in wild type but slower than in the isolated γDC$_{NTD}$. C) Subtractive deuteration plot of dInt-γDC and V132A γDC.

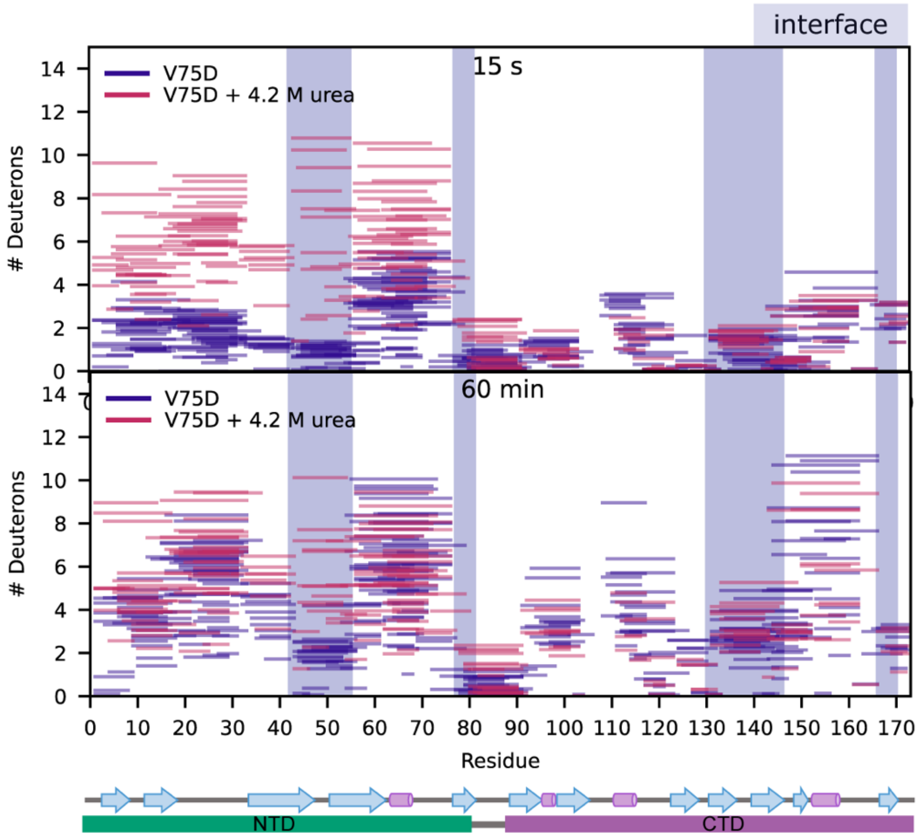


**Figure S3: Comparison of V75D HDX with and without urea.** Number of deuterons uptaken by V75D under native conditions (blue) and in 4.2 M urea (magenta) at 15 seconds (top) or 60 minutes (bottom). Each line represents an individual peptide spanning the residues indicated on the x-axis, with number of deuterons uptaken indicated on the y-axis. After 60 minutes, exchange profiles of V75D under native conditions mostly match V75D in urea, with notable exceptions at the NTD interface (interfacial regions are highlighted in pale blue). Secondary structure map and domain regions are indicated below the plot.

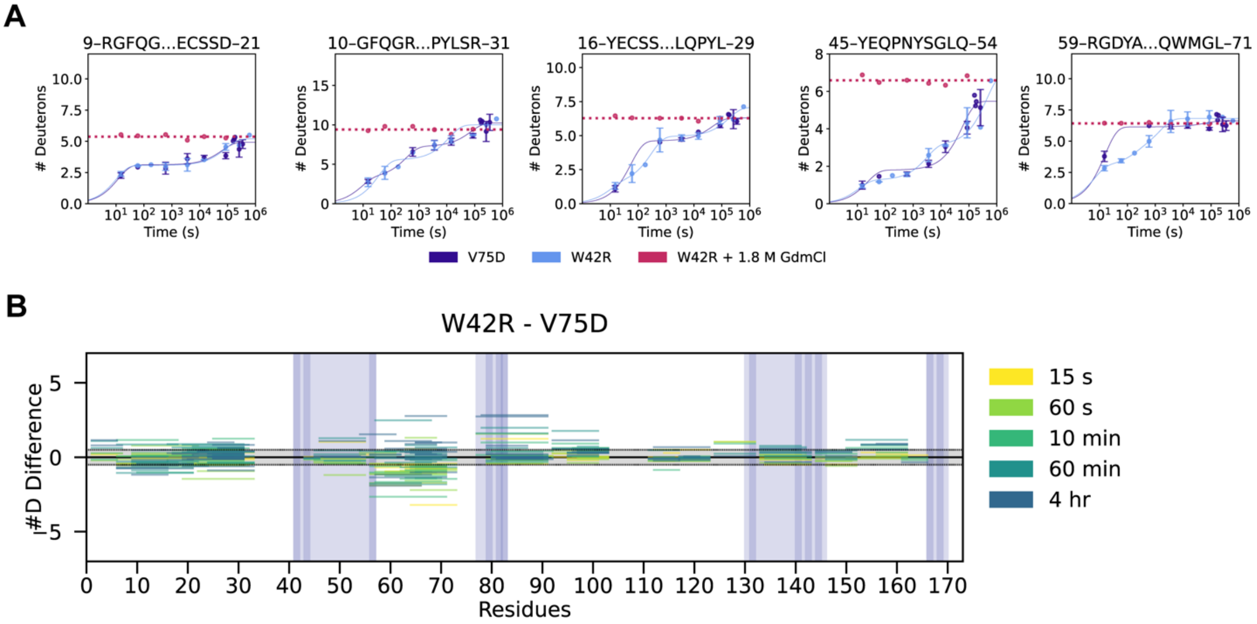


**Figure S4: W42R γDC is comparable to V75D γDC in exchange behavior.** A) Uptake plots comparing W42R, V75D, and W42R + 1.8 M GdmCl. The dotted magenta line is average deuteron uptake of W42R in 1.8 M GdmCl. B) Subtractive deuteration plot of W42R and V75D.

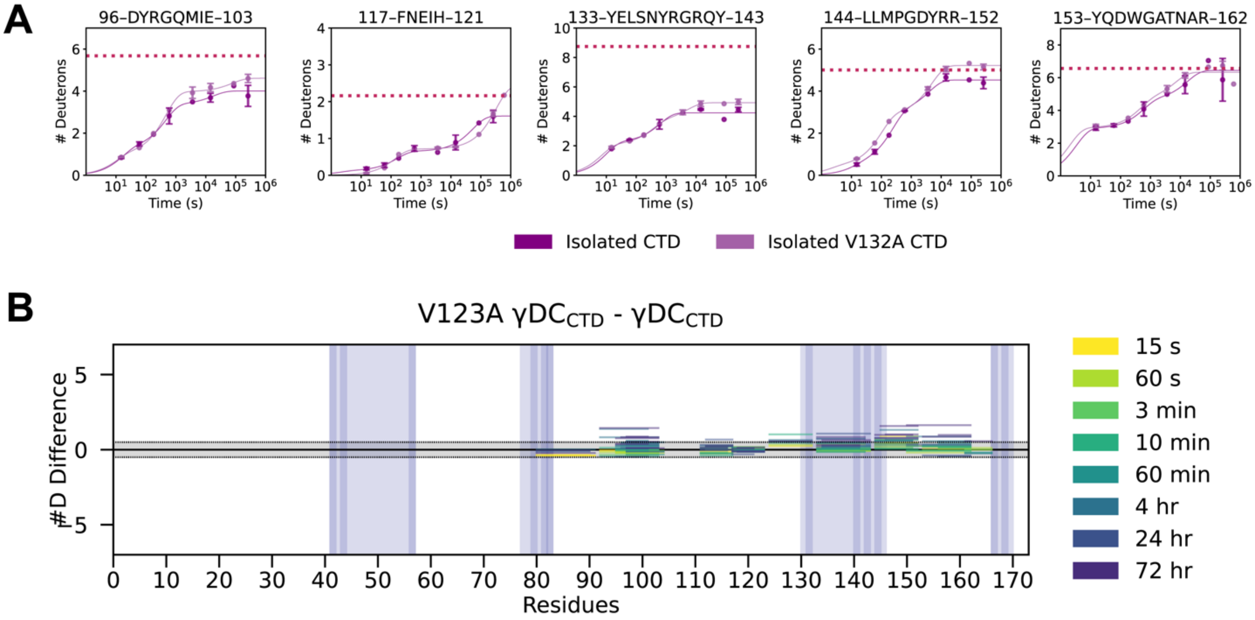


**Figure S5: V132A has only a mild impact on exchange of γDC$_{CTD}$.** A) Uptake plots depicting V132A γDC$_{CTD}$ and wild-type γDC$_{CTD}$. The dotted magenta line is average deuteron uptake of V132A in 4.8 M GdmCl. B) Subtractive deuteration plot of V132A γDC$_{CTD}$ and wild-type γDC$_{CTD}$. V132A γDC$_{CTD}$ has minor loss of protection in peptides, especially near position 132, but otherwise closely resembles the wild-type γDC$_{CTD}$.

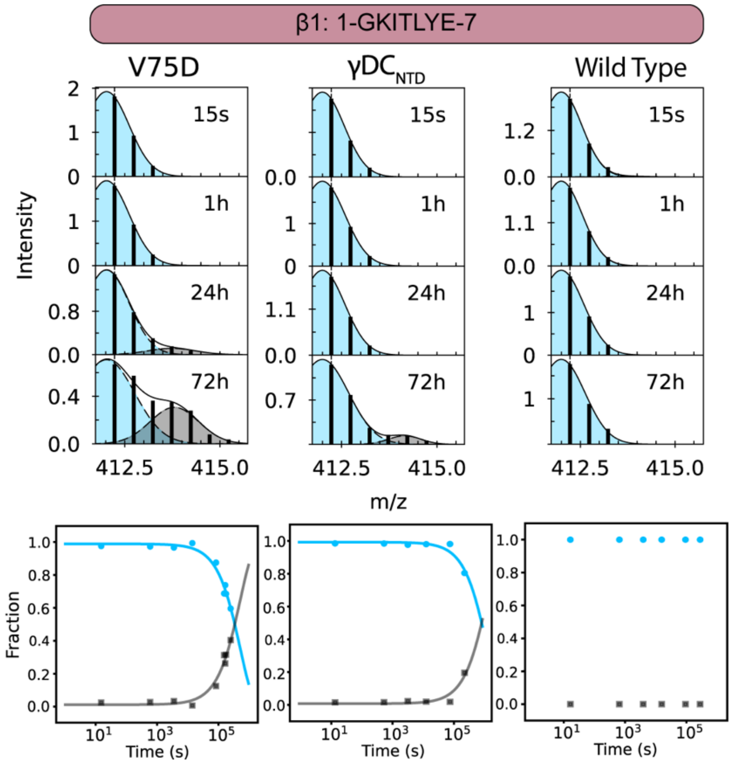


**Figure S6: β1 EX1 exchange mass spectra.** Top: Representative mass spectra of a β1 peptide in V75D, γDC$_{NTD}$, and wild-type as time of exchange increases, with overlaid Gaussian fits to indicate two populations (light blue, lighter, less-exchanged population; gray, heavier, more-exchanged population). Bottom: EX1 kinetics of conversion from light to heavy (wild type, too slow to fit).

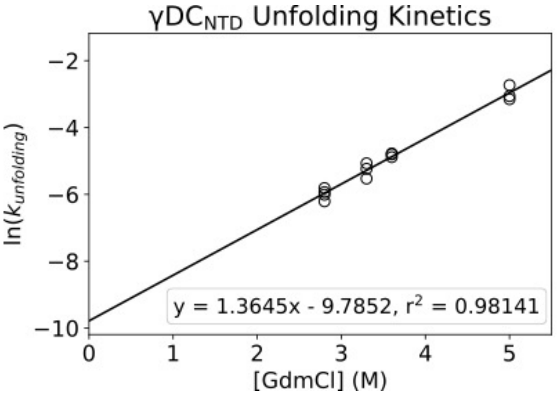


**Figure S7: Unfolding rate of γDC$_{NTD}$ in GdmCl.** The natural log of the observed unfolding rate of γDC$_{NTD}$ in GdmCl as measured by increase in fluorescence at 350 nm (excitation 280 nm) is linear with respect to denaturant concentration. Linear extrapolation to 0 M denaturant predicts an unfolding rate of 5.6 x $10^{-5}$ $s^{-1}$, which corresponds to a folding rate of ~ 0.47 $s^{-1}$ (all assuming two-state unfolding behavior).

**Table S1: Bootstrapped parameters for Ising fits.** Equilibrium two-state, three-state, and global Ising fit parameters for all γDC variants. ΔG˚ (folding) values are in kcal/mol; *m*-values are in kcal $mol^{-1}$ $M^{-1}$ GdmCl). Values are from 3,000 bootstrapping iterations. For V132A, $\Delta G^{\circ}_{interface}$ was held equivalent to $\Delta G^{\circ}_{interface\ (NTD\ unfolded)}$. Abbreviations: RSSR, reduced sum of square residuals; CI, confidence interval.
[a] Value obtained from a separate fit incorporating this term (no significant change to other parameters).

| **Wild-type γDC** | | | |
|---|---|---|---|
| RSSR = $4.11 \times 10^{-4}$ | Mean | 5% CI | 95% CI |
| $\Delta G^{\circ}_{NTD}$ | -4.48 | -4.89 | -4.08 |
| $\Delta G^{\circ}_{CTD}$ | -8.24 | -8.97 | -7.59 |
| $\Delta G^{\circ}_{interface}$ | -3.31 | -4.03 | -2.52 |
| $\Delta G^{\circ}_{interface\ (NTD\ unfolded)}$[a] | 0.02 | -0.26 | 0.27 |
| $m_{NTD}$ | 3.15 | 2.87 | 3.44 |
| $m_{CTD}$ | 2.61 | 2.41 | 2.85 |
| **V132A γDC** | | | |
| RSSR = $5.12 \times 10^{-4}$ | Mean | 5% CI | 95% CI |
| $\Delta G^{\circ}_{NTD}$ | -4.54 | -5.09 | -4.04 |
| $\Delta G^{\circ}_{CTD}$ | -6.74 | -7.47 | -6.11 |
| $\Delta G^{\circ}_{interface}$ | -1.21 | -1.52 | -0.92 |
| $\Delta G^{\circ}_{interface\ (NTD\ unfolded)}$ | -1.21 | -1.52 | -0.92 |
| $m_{NTD}$ | 3.20 | 2.84 | 3.57 |
| $m_{CTD}$ | 2.88 | 2.61 | 3.18 |

**Table S2:** Average redundancy, coverage, and back exchange as well as number of replicates per each construct studied with HDX-MS. Replicates given as number of technical replicates collected. Average back exchange calculated from angiotensin-II as 100% - #D/(5*90%) (angiotensin-II has five exchangeable protons). Coverage given as percentage of residues contained in at least one peptide. Redundancy given as average number of peptides per amide. Repeatability given as average standard deviation of deuteration of all peptides across replicates (Da). Significant differences in HDX given as 95% confidence intervals (Da).

| | **V75D** | **γDC$_{NTD}$** | **γDC$_{CTD}$** | **V132A** | **V132A γDC$_{CTD}$** | **V75D/ V132A** | **Wild-type γDC** | **W42R** | **dInt-γDC** |
|---|---|---|---|---|---|---|---|---|---|
| **Replicates** | 5 | 2 | 2 | 3 | 2 | 2 | 2 | 2 | 1 |
| **Back exchange** | 34.0% | 29.7% | 28.2% | 30.4% | 28.6% | 25.9% | 26.4% | 29.2% | 25.3% |
| **Coverage** | 90.6% | 96.3% | 84.2% | 79.6% | 75.3% | 83.2% | 86.1% | 89.3% | 87.9% |
| **Redundancy** | 10.6 | 23.1 | 4.2 | 7.5 | 2.7 | 5.7 | 12.1 | 10.9 | 9.9 |
| **Avg. peptide length** | 11.1 | 12.4 | 8.5 | 11.9 | 7.9 | 10.1 | 11.1 | 11.0 | 11.8 |
| **Avg. # peptides** | 192.4 | 176.5 | 58 | 132 | 42.5 | 120.5 | 212 | 210 | 175 |
| **Repeatability** | 0.31 | 0.23 | 0.26 | 0.22 | 0.11 | 0.22 | 0.28 | 0.39 | |
| **Significant differences in HDX (95% CI)** | 0.37 | 0.34 | 0.39 | 0.27 | 0.16 | 0.33 | 0.41 | 0.61 | |

**Table S3: Details of Ising partition functions and fraction folded expressions used for global 1-D Ising analysis of wild-type and V132A γDC.** The equilibrium constant $\kappa$ for an individual domain *i* as a function of GdmCl is $\kappa_i = e^{-(\Delta G_i + m_i x)/(RT)}$, and the equilibrium constant $\tau$ for the interface coupling between the NTD and the CTD is $\tau_{N,C} = e^{-\Delta G_{N,C}/(RT)}$. The equilibrium constant $\omega$ describing coupling between the unfolded NTD and the folded CTD is $\omega_C = e^{-\Delta G_{unfoldedN,C}/(RT)}$. For V132A, $\omega_C$ was held equivalent to $\tau_{N,C}$.

| | | **1-D Ising Model** | **Modified 1-D Ising Model** |
|---|---|---|---|
| **Isolated domain model (two-state)** | Partition function | $\rho = 1 + \kappa_i$ | $\rho = 1 + \kappa_i$ |
| **Isolated domain model (two-state)** | Fraction folded | $\frac{\kappa_i}{\rho}$ | $\frac{\kappa_i}{\rho}$ |
| **Full construct model (four-state)** | Partition function | $\rho = 1 + \kappa_N + \kappa_C + \kappa_N \kappa_C \tau_{N,C}$ | $\rho = 1 + \kappa_N + \omega_C \kappa_C + \kappa_N \kappa_C \tau_{N,C}$ |
| **Full construct model (four-state)** | Fraction unfolded | $1 - \frac{\kappa_N \kappa_C \tau_{N,C} + \kappa_C/2 + \kappa_N/2}{\rho}$ | $\frac{1}{\rho}$ |
| **Full construct model (four-state)** | Fraction partially folded | -- | $\frac{\omega_C \kappa_C + \kappa_N}{\rho}$ |
| **Full construct model (four-state)** | Fraction fully folded | $\frac{\kappa_N \kappa_C \tau_{N,C} + \kappa_C/2 + \kappa_N/2}{\rho}$ | $\frac{\kappa_N \kappa_C \tau_{N,C}}{\rho}$ |
